# Centriolar subdistal appendages promote double strand break repair through homologous recombination

**DOI:** 10.1101/2022.10.19.512819

**Authors:** Guillermo Rodríguez-Real, Andrés Domínguez-Calvo, Rosario Prados-Carvajal, Aleix Bayona-Feliú, Sónia Gomes-Pereira, Fernando R. Balestra, Pablo Huertas

**Author notes:** Lead Contact: Pablo Huertas.

## Abstract

The centrosome is a cytoplasmic organelle with roles in microtubule organization which has also been proposed to act as a hub for cellular signaling. Some centrosomal components are required for full activation of the DNA Damage Response. However, if the centrosome regulates specific DNA repair pathways is not known. Here, we show that centrosomes presence is required to fully activate recombination, specifically to completely license its initial step, the so-called DNA end resection. Furthermore, we identify a centriolar structure, the subdistal appendages, and a specific factor, CEP170, as the critical centrosomal component involved in the regulation of recombination and resection, albeit it does not control end-joining repair. Cells lacking centrosomes or depleted for CEP170 are, consequently, hyper-sensitive to DNA damaging agents. Moreover, low levels of CEP170 in multiple cancer types correlate with an increase of the mutation burden associated with specific mutational signatures and a better prognosis, suggesting that changes in CEP170 can act as a mutation driver but also could be targeted to improve current oncological treatments.

## Introduction

The centrosome is considered the major microtubule (MT) organizing center (MTOC) of animal cells, involved in mitotic spindle assembly, interphase MT-network organization and cilia and flagella assembly. Centrosomal defects, either in number or structure, are linked to chromosome missegregation, polarity defects and motility or signaling defects related to the lack of cilia or flagella (Bornens, 2012; Nigg & Raff, 2009). The centrosome is composed of a pair of centrioles surrounded by a structured cloud of proteins known as the pericentriolar material (PCM) (Le Guennec *et al*, 2021). The centriolar structure and its assembly pathway has been conserved throughout evolution (Nabais *et al*, 2020). In humans, they are composed by 9 triplets of MTs distributed in a radial symmetry to form a cylinder of approximately 500 nm long and 200 nm wide. In addition to MTs, from two to three hundred proteins have been estimated to localize to the centrosome (Andersen *et al*, 2003; Jakobsen *et al*, 2011). However, only a minority of them has been described to play a role at the centrosome. For instance, the cartwheel component HsSAS-6 or the kinase PLK4 are required for centriolar duplication, thus the inactivation or depletion of either protein in human cultured cells leads to the inhibition of centriole assembly, rendering the appearance of cells without centrioles as the subsequent divisions go by (Wong *et al*, 2015; Wang *et al*, 2015; Gönczy & Hatzopoulos, 2019). Interestingly, centriole depleted cells can assemble a mitotic spindle and perform mitosis, although with a high rate of chromosome missegregation (Wong *et al*, 2015; Wang *et al*, 2015). These cells without centrioles expend a significant longer time in mitosis which leads to the activation of a p53-dependent pathway that leads to a G1 cell cycle arrest (Meitinger *et al*, 2016; Wong *et al*, 2015; Lambrus *et al*, 2016; Fong *et al*, 2016; Lambrus *et al*, 2015). Other well characterized centriolar proteins are the components of the centriolar appendages, substructures present only in mature centrioles (i.e. those that are at least two cell cycle old). Centriolar appendages protrudes from the distal side of the centriole usually displaying a nine-fold radial symmetry and they are classified into two types depending on their position within the centriole: the distal and the subdistal appendages (Le Guennec *et al*, 2021). Centriolar proteins required to build these structures have been identified including ODF2, CEP128, Centriolin, NDEL1, Ninein and CEP170 as key components required to build the subdistal appendages. From a functional point of view, among other roles, distal appendages are essential for cilia assembly and subdistal appendages are involved in MTs anchoring to the centriole (Hall & Hehnly, 2021).

Interestingly, it has been proposed that the centrosome plays also cellular functions not directly related to its MTOC activity. One of these functions involves its role as a signaling platform to regulate several cellular processes such as cell polarization, cell cycle regulation or cell proliferation (Arquint *et al*, 2014). Intriguingly, several components of the DNA damage response (DDR) have been identified at the centrosome as the checkpoint proteins ATM, ATR, Chk1 or Chk2 and the repair factors BRCA1, BRCA2 or PARPs (Mullee & Morrison, 2016). Additionally, several centrosomal proteins have been shown to play a direct role in specific DNA damage repair pathways, for example Centrin-2 in nucleotide excision repair (Dantas *et al*, 2011), or PCNT and CEP164 in checkpoint triggering by ATR and ATM activation (Sivasubramaniam *et al*, 2008; Griffith *et al*, 2008), although the role of CEP164 in DNA repair has been recently disputed (Daly *et al*, 2016). However, it remains unclear if and how the centrosome itself plays a role specifically in the regulation of the DNA double strand breaks (DSBs) repair pathways.

DSBs are one of the most challenging DNA lesions to be fixed. As the rest of DNA damage, they can arise due to exposure to exogenous (chemicals, irradiation, etc.) or endogenous (transcription or replication problems) (Ciccia & Elledge, 2010; Tubbs & Nussenzweig, 2017; Jackson & Bartek, 2009). In contrast to other DNA lesions, DSBs lack an intact DNA strand to use as a template during repair, hence they are particularly complex to repair. Indeed, DSBs appearance triggers a complex response, the already mentioned DDR, which overhauls the whole cellular metabolism. The DDR is triggered mainly by the activation of the kinases ATM, ATR, Chk1 and Chk2 (Blackford & Jackson, 2017; Ciccia & Elledge, 2010), in a complex process that, as mentioned, requires partially centrosomal components (Mullee & Morrison, 2016). Furthermore, DSBs could be repaired by using one out of several possible mechanisms that can be grouped in two major families: the homologous recombination pathways (HR) or the non-homologous end-joining pathway (NHEJ) (Chang *et al*, 2017; Ranjha *et al*, 2018; Jasin & Rothstein, 2013). Upon the detection of a DSB, the cell needs to commit to one of these two alternative repair pathways, and the outcome of the repair in terms of fidelity is dependent on this decision. The choice is influenced by different cellular cues, such as cell cycle stage. HR can almost only be executed after DNA replication (in S and G2 phases) when the sister chromatid is present and can be used as a template to repair the damaged molecule (Symington, 2016; Jasin & Rothstein, 2013). On the contrary, NHEJ that relies on the ligation of the two ends of DNA with little processing and without a template, can be performed through the whole cell cycle (Chang *et al*, 2017). A complex molecular network is involved in determining the repair pathway of choice (HR vs. NHEJ) upon the detection of a DSB, thus affecting DSB repair outcome. Nevertheless, it happens mostly by controlling the initial step of HR, the so-called DNA end resection (Cejka, 2015; Symington, 2016). This consists in the extensive processing of the DNA ends by nucleases to form long tails of 3’ OH protruding long tails of single stranded DNA (ssDNA). These ssDNAs are initially coated by the RPA complex (RPA1, RPA2, and RPA3), which protects it from degradation and from pairing with other ssDNAs. Importantly, resected DNA effectively blocks NHEJ, hence committing the repair through HR. Mostly, cellular cues that promote the licensing of DNA end resection, thus promoting HR, affect the activation of the nuclease activity of the MRN complex (MRE11-RAD50-NBS1) through the regulation of CtIP and its partner BRCA1 (Symington, 2016; Cejka, 2015). This triggers the so-called short-range DNA end resection that is followed by a more extensive resection, known as long-range resection (Symington, 2016; Cejka, 2015). From that point onward, HR can happen by different means in a complex molecular choreography that ends with the use of a homologous template for repair (Jasin & Rothstein, 2013). Among other factors, it usually requires the recruitment of the recombinase Rad51 that modulates the invasion of the homologous template.

Interestingly, among the proteins identified to localize to the centrosome, in addition of proteins involved in the DDR (such as ATM, ATR, Chk1 or Chk2) there is a subgroup of them linked directly to DSBs repair by HR (i.e. BRCA1 and BRCA2). Thus, we wondered whether the centrosome might play a direct role in HR and, more specifically, in regulating the choice between this repair mechanism and NHEJ by affecting DNA end resection. Strikingly, we observed that cells without centrioles are defective in DNA end resection and HR, supporting this hypothesis. Moreover, such regulation of DNA end resection at centrioles relies on the subdistal appendages. We found that phosphorylation of the subdistal appendage protein CEP170 by the checkpoint kinases ATM and ATR is required to fully activate HR pathway. In the absence of centrioles, subdistal appendages or CEP170, HR is defective, rendering cells hyper-sensitive to DNA damaging agents. Strikingly, CEP170 levels are altered in several tumors. Interestingly, low levels of the protein increase their mutational burden with specific mutational signatures, chiefly among them the one associated with HR defect. Furthermore, low levels of CEP170 levels correlates with poorer cancer prognosis.

## Results

### The centrosome promotes the repair of DNA Double Strand Breaks through homologous recombination

Despite the aforementioned connection between the centrosome and the DDR, it is unknown if there is a direct centrosomal role on DSB repair pathway choice. Intrigued by this possibility, we generated U2OS cells without centrosomes using the PLK4 inhibitor centrinone (Wong *et al*, 2015). As mentioned, PLK4 activity is required for centriole duplication, thus its inhibition leads to cells that while dividing in subsequent cell cycles dilute their preexisting centrioles rendering to the appearance of centrosome-less cells in the population. Centriolar loss requires therefore several cell divisions, hence cannot be observed upon short treatments but only after exposure to the drug for several days (see Figure EV1A and B comparing the 1 day *vs* 7 days time points). We then used this condition to test the impact of centriole loss on the balance between NHEJ and HR repair pathway choice (Figure 1A-B). We used a duo of stably transfected GFP-based repair reporters in which the ectopic expression of the I-SceI nuclease fused to BFP (Blue fluorescent protein) triggers the reconstitution of an active GFP when a specific repair pathway, either HR or NHEJ, is activated (Pierce *et al*, 1999; Bennardo *et al*, 2008) (Figure 1A-B). In this experimental set up, infection efficiency for all conditions were monitored and only BFP positive cells were taken into consideration for GFP analysis (for more details see Material and Methods). Strikingly, centriole depleted cells, i.e. treated with centrinone for 7 days, compromised the homologous recombination Rad51-dependent gene conversion pathway (Figure 1A) and at the same time promoted the error prone NHEJ repair (Figure 1B), suggesting that indeed centrosomes are critical to balance DSB-repair pathway choice. As introduced, this decision occurs usually by regulating HR, and mostly by licensing or not DNA end resection. To dissect at which step the centrosome impinges on this process, we monitored by immunofluorescence key steps of the homologous recombination repair pathway. Specifically, we inspected the steps of DNA end resection (RPA foci), of HR initiation/commitment (BRCA1 foci), and strand invasion for HR (RAD51 foci) in U2OS cells depleted for centrioles and challenged with 10Gy of ionizing radiation. Centrosome depleted cells showed a significant drop in all three steps of HR (Figure 1C-E) suggesting that centrosomes are required to activate HR from the earliest step, i.e. affecting already the licensing of DNA end resection. Similar resection impairment was observed in centrinone treated U2OS cells when DNA damage was induced by Camptothecin (Figure EV1C) or in centrinone treated RPE-1 cells irradiated with 10 Gy (Figure EV1D). To discard that centrinone itself could directly have an effect in HR independently of its effect on centriole duplication, we repeated the RPA foci formation experiment in cells treated with centrinone but only for one day, thus in which the majority of cells retain centrioles, and no impact on resection was observed (Figure EV1E). Finally, to confirm that centriole loss and not extended centrinone treatment by other means impair HR, we repeated the same experimental strategy making use of the HsSAS-6 knock out RPE-1 p53-/- cell line. HsSAS-6 is an essential protein for centriole duplication and therefore HsSAS-6 KO cells are devoid of centrosomes and they are only viable in the absence of p53 (Wang *et al*, 2015; Leidel *et al*, 2005). In agreement with our previous results, HsSAS-6 KO p53 -/- cells also showed a significant decrease in resection compared to control RPE-1 p53 -/- cells (Figure EV1F).

**Figure 1:**
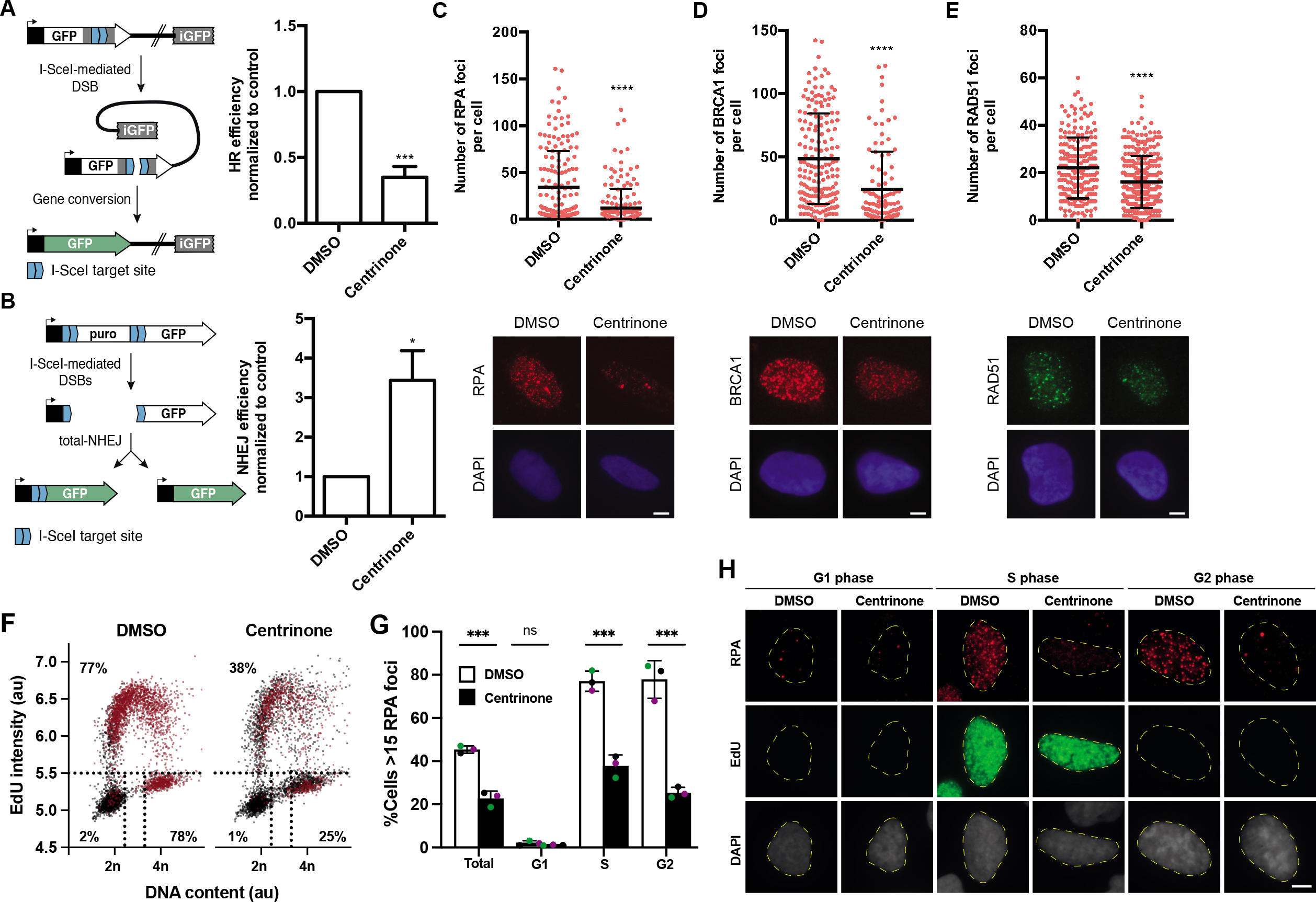
Centrosomes control the balance between repair mechanisms. **A**, Homologous recombination was assessed in U2OS cells bearing the DR-GFP reporter in cells lacking centrosomes upon the treatment with centrinone for 7 days or mock-treated with DMSO. A schematic representation of the reporter and the recombination event that renders the appearance of a functional GFP gene is shown on the left side. Within the BFP positive cell population, i.e. cells transfected with I-Sce-I containing viral particles, the percentage of GFP positive cells, i.e. cells that have undergone repair by HR after induction with the nuclease I-SceI, was normalized to the DMSO control taken as 1. **B**, Same as A but in cells harboring the NHEJ reporter EJ5-GFP. **C**, U2OS cells treated for 7 days with centrinone or DMSO, as indicated, were irradiated with 10 Gy and 1 hour later prepared for immunofluorescence using an anti-RPA antibody as described in the methods section. The number of RPA foci per cell for at least 200 cells per condition was quantified automatically using FIJI software and plotted. One representative experiment out of three performed with similar results is shown. (top). Representative images are shown (bottom). Scale bar in white represents 7 μm. **D**, Same as C, but using an antibody against BRCA1. **E**, Same as C, but using an anti-RAD51 antibody. **F**, U2OS cells were treated 7 days with centrinone or DMSO as control before a 30-minutes pulse-labeled with EdU. Afterward, cells were irradiated (10 Gy) and incubated for one hour. Click-it reaction and immunostaining against RPA was carried out as described in the methods section. Automated multichannel wide-field microscopy for quantitative image-based cytometry (QIBC) was performed. Single-cell data for more than 5000 cells is depicted as scatter plots showing DNA content (DAPI intensity) and EdU incorporation during S phase (EdU intensity). Correlatively, RPA foci was measured for each cell. Cells with more than 15 RPA foci are classified as positive (labelled in red). **G**, Quantification of RPA foci positive cells regarding cell cycle status from three different experiments analyzed as described in F. **H**, Representative images of F and G are shown. Scale bar in white represents 7 μm. In A, B and G the average and standard deviation of three independent experiments is shown. The statistical significance was calculated using a Student’s t-test. p values are represented with one (p < 0.05), two (p < 0.01), three (p < 0.001) or four (p < 0.0001) asterisks. Non statistical significance is labeled ns.

Since all our data supported a role of the centrosome as a regulator of HR, we wondered whether this regulation relied on its presence/absence or the number of centrosomes. Thus, we wondered if cells bearing an excess number of centrosomes could also display an imbalance in HR *vs.* NHEJ. To test this idea, we transiently overexpressed an mCherry-tagged version of PLK4 (Moyer *et al*, 2015) which leads to ∼35% of cells bearing extra centrosomes in our cell population (Figure EV1G). Notably, we could not observe any imbalance between HR *vs.* NHEJ in this condition (Figure EV1H and I) suggesting that presence or absence, but not number of centrosomes regulates double strand break repair pathway choice.

We wonder if centrosome lost itself could induce DNA double strand breaks. To test this idea, we used the phosphorylated form of the histone variant H2AX (γH2AX) as a marker of DSB. As expected, we did not observe any impact on γH2AX foci formation upon centrinone treatment (Figure EV1J). We then checked whether the imbalance observed in centrosome depleted cells between the two main double strand break repair pathways would eventually lead to a delay or a lack of repair or if it will simply result on a switch between HR and NHEJ. To do so, we compared the foci dynamics of γH2AX for 6 hours after irradiation in control and centriole depleted cells (Figure EV1K). Notably, we did not observe any significant change in the dynamics of chromosome break repair in centriole depleted cells, supporting the idea that the decrease of double strand break repair by HR is countervailed by an increase in NHEJ repair pathway (Fig. 1A and 1B). Intriguingly, we did not observe an overall increase on NHEJ regulators, such as RIF1 nuclear foci, in irradiated centriole depleted cells (Figure EV1L). If any, we observed a slight reduction on RIF1 accumulation. As RIF1 is an accessory NHEJ factor involved in counteracting resection, our data argue that the increase in NHEJ observed in acentriolar cells does not simply rely on hampering DNA end processing. We then wondered about the mechanism by which the centrosome could be regulating HR *vs*. NHEJ repair pathway choice. We hypothesized that the role of the centrosome in DSB repair might rely on its activity as a microtubule nucleator during interphase. To test this, we measured resection in irradiated U2OS cells pretreated either with MTs depolymerizing (Nocodazole) or stabilizing (Taxol) drugs (Figure EV1M). Neither of the two treatments mimicked the impact of centrosome loss, suggesting that the connection of the centrosome with DSB repair is independent of its MTOC activity. Overall, we concluded that by a so far uncharacterized mechanism, the centrosome promotes DNA DSB repair by stimulating homologous recombination and impairing NHEJ in a fashion that is independent of its MT organization function.

### DNA double strand break repair imbalance upon centrosome loss is not a consequence of cell cycle perturbance

Notably, extended centrinone treatment has been reported to prolong mitosis through a 53BP1-USP28-mediated activation of p53 and p21, which eventually blocks the cell cycle in G1 (Meitinger *et al*, 2016; Wong *et al*, 2015; Fong *et al*, 2016; Lambrus *et al*, 2016). As cell cycle is a major regulator of DNA end resection and HR (Huertas, 2010), we wondered if such response could account for the imbalance observed in DNA double strand break repair. Interestingly, in our experimental conditions, we did not observe either a significant accumulation in G1, an increase of mitotic cells or cell death in cells treated with centrinone (Figure EV1N-Q). Moreover, our results in HsSAS-6 KO cells were already obtained in a p53 KO background, reinforcing the idea that this effect was cell cycle independent (Figure EV1F). Despite that, to further rule out that the absence of centrosomes could be modulating HR indirectly through changes in cell cycle progression mediated by p53, we performed similar experiments in not only Saos-2 cells, a primary osteosarcoma p53-/- cell line, but also in RPE-1 lacking either p53 or p21 (Hauge *et al*, 2019). We obtained similar results to the ones observed in U2OS and RPE-1 cells with all cell lines (Figure EV1R-Z), supporting the idea that the effect of centriolar loss on HR is independent of an accumulation in G1 or mitosis. However, HR takes place during S and G2, and centrinone treatment increases the number of cells in G2 (Figure EV1N). Although we already observed that treatment with camptothecin, that specifically creates DSBs at S phase, was compromised upon centrinone treatment (Figure EV1C), we tested if centriole loss affected equally DNA end resection in these both phases of the cell cycle by using quantitative image-based cytometry (QIBC) upon irradiation (Figure 1F-H). With this approach, we measured DNA content (DAPI intensity), active DNA replication (with an EdU pulse) and DNA resection (through RPA foci), by fluorescent microscopy (Figure 1F-H). As expected, no resection was observed in G1 cells in any condition. Furthermore, this analysis confirmed that the decrease in resection upon centrinone treatment is similar in both S and G2 phases.

### Centriolar subdistal appendages proteins promote DNA end resection after double strand break

As stated in the introduction, a few DDR and HR proteins are located at the centrosome (ATM, ATR, CHK1, CHK2, BRCA1, BRCA2) and some centriolar proteins have been previously involved in DDR (Centrin2, CEP63, PCNT) (Mullee & Morrison, 2016; Dantas *et al*, 2011; Sivasubramaniam *et al*, 2008; Alderton *et al*, 2006), suggesting a close crosstalk between them. We envisaged that the regulation of DSB repair pathway choice by centrosomes will require one or several proteins sharing either of these characteristics: a DDR protein that resides temporarily at the centrosome or a centrosomal-factor that regulates and/or is regulated by the DDR. To search for these factors with an unbiased approach we decided to retrieve previously published data from several publications and databases of either DNA repair/DDR/DDR-regulated factors or centriole/centrosome associated proteins and performed an *in silico* metanalysis to look for common factors within these two categories. This metanalysis retrieved a list of genes previously connected to DDR (1457 genes) and to centrosome biology (1424 genes). The cross-check of these two lists led to the identification of 159 genes linked to both processes (Figure 2A; Figure EV2A; Table EV1). Among these 159 genes, caught our attention a group of 3 genes previously reported to be part of a very defined substructure of the centrioles, the subdistal appendages (CEP128, NIN and CEP170) (Hall & Hehnly, 2021). A handful of proteins have been identified to be part of such structure: ODF2, CEP128, Centriolin, NDEL1, Ninein and CEP170. Their architecture follows a hierarchical organization with ODF2 at the root, and the most proximal to the centriole wall. From that point the structure branches, with a short bough formed by NDEL1 and a second longer one formed by Cep128-Centriolin-Ninein and CEP170, in that order from the inner part to the periphery (Figure 2B). To assess whether the subdistal appendages as a structure is involved in DSB repair, and most specifically in activating HR by licensing DNA end resection, we depleted the above-mentioned proteins using siRNA against them. Depletion of these components was confirmed either by western blot or by RT-PCR, depending on the availability of specific antibodies (Figure EV2B-D). Interestingly, while reduction of ODF2, Centriolin, Ninein and CEP170 showed a significant drop in DNA end resection after DNA damage, albeit to different extents, depletion of NDEL1 did not have a significant impact on the process (Figure 2C). Knockdown of the core resection factor CtIP was used as a positive control. Thus, we conclude that the subdistal appendages, at least the longer branch, is required for fully activation of DNA end resection. Intrigued by these results, we reasoned that the most peripheral component of the long branch of the subdistal appendages, CEP170, could be the critical factor involved in DNA end processing. Therefore, the effect of the depletion of other subdistal appendages proteins might reflect their impact on CEP170 localization to this centriolar structure. To test this idea, we quantified CEP170 levels at the centrosome after depletion of all the subdistal appendages proteins using the same samples that were used for RPA foci formation (Figure 2D). As published, depletion of ODF2, CEP128, Centriolin and Ninein, but not NDEL1, affected CEP170 localization at centrioles to different extents. As expected, downregulation of the core resection factor CtIP did not affect this localization either. Strikingly, and in agreement with our hypothesis, we found a correlation between the decrease of CEP170 levels at the centrosome and the drop on DNA end resection (Figure 2E; R^2^=0.9799). Furthermore, this drop in DNA end resection was also confirmed in the RPE-1 CEP128 KO cell line (Mönnich *et al*, 2018) where CEP170 levels at the centrosome is also partially reduced (Figure EV2E-F). CEP170 depletion phenotype was rescued by a siRNA-resistant version of the gene (Figure 2F and Figure EV2G). Moreover, the drop in DNA end resection after CEP170 depletion was also observed when cells were challenged with the DNA damaging agent Camptothecin (Figure EV2H). We also confirmed that CEP170 depletion itself has no impact on RPA or γH2AX foci formation in the absence of an exogenous source DNA damage (Figure EV2I-J). Lastly, we used CRISPR/Cas9 technique to create a KO of CEP170 in RPE cells. Unfortunately, we were not able to recover a full KO, suggesting the protein is essential. However, we created several independent heterozygous CEP170 +/- modified cell lines (Figure EV2K). All those heterozygous cell lines for CEP170 also displayed a clear drop in DNA end resection (Figure 2G). All together, these results revealed that the subdistal appendages are key structures within the centriole involved in DNA end resection and point to CEP170 as the critical mediator of this process.

**Figure 2:**
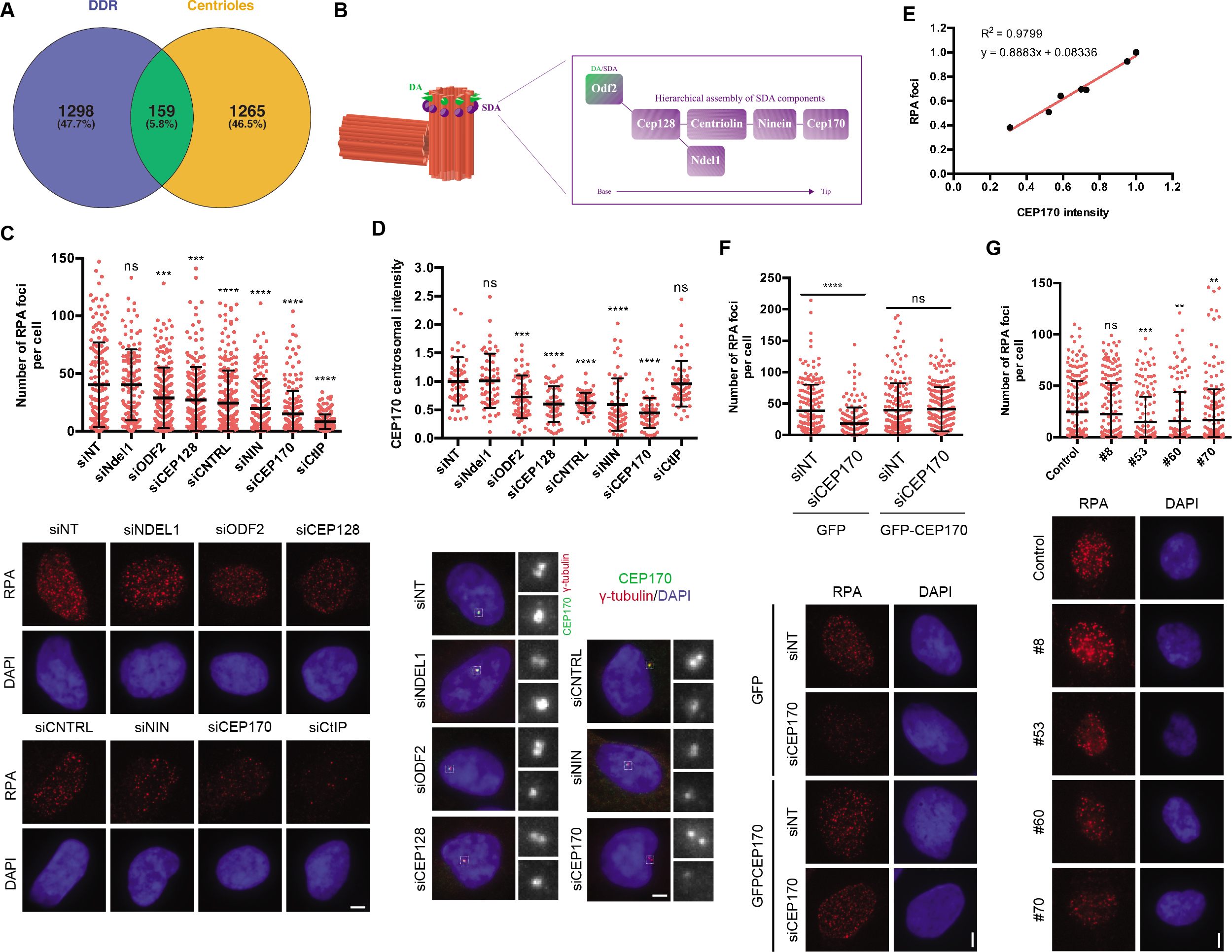
CEP170 and the Subdistal appendages regulate DNA end resection. **A**, Venn diagram showing the overlapping of genes found both in DDR screenings and centrosome related screening, as described in the main text and the method section. **B**, Schematics of the centrioles showing the Distal and Subdistal appendages (DA and SDA, respectively). The inset shows the molecular architecture of the SDA. **C**, Cells transfected with siRNAs against the indicated proteins or a siRNA control (siNT) and the number of RPA foci per cell for at least 200 cells per condition was quantified automatically using FIJI software and plotted. One representative experiment out of three performed with similar results is shown (top). Representative images are shown (bottom). Scale bar in white represents 7 μm. **D**, the same samples used in C were used to immunodetect CEP170 at centrioles. Intensity of individual cells is plotted (top) in cells depleted for the indicated factors. Representative images of each condition are shown on the bottom side. For each condition, insets of centrosomes labelled by either CEP170 or γ-tubulin are shown. **E**, linear correlation between RPA foci and CEP170 intensity. For each condition, the average calculated in C and D was used in the plot. The R^2^, linear correlation and mathematical formula are shown. **F**, cells harboring a plasmid expressing an siRNA-resistant GFP-tagged CEP170 or an empty vector were transfected with siRNAs against CEP170 or a control sequence (siNT). Then, cells were treated with IR and the number of RPA foci per cell was quantified as in C (top). Representative images are shown (bottom). **G**, Same as C but in RPE-1 cells with partial knock out of CEP170 (top). Representative images are shown (bottom). In C, D, F, and G, the statistical significance was calculated using a Student’s t-test. p values are represented with one (p < 0.05), two (p < 0.01), three (p < 0.001) or four (p < 0.0001) asterisks. Non statistical significance is labeled ns.

### CEP170 promotes homologous recombination from the centrosome

Our data suggest that the subdistal appendage component CEP170 promotes DNA end resection after DSB damage, and this DNA processing requires the presence of centrioles. To link those two observations, and test if CEP170 was affecting resection from the centrosome and not due to a putative, unknown, centrosome-independent function, we tested the effect of CEP170 depletion on DNA end resection in cells without centrioles. As control, we performed similar experiments depleting the HR regulator CtIP. Interestingly, we observed that while CtIP depletion in cells without centrioles further reduced DNA end resection, depletion of CEP170 in cells without centrioles did not show any impact (Figure 3A), thus suggesting that the loss of centrioles is epistatic over CEP170 role but not over a core resection protein such as CtIP. Therefore, the lack of an additive phenotype in centriole-less cells depleted of CEP170 strongly support that CEP170’s role in DDR is played from the centrosome. Furthermore, as the absence of centrioles and the depletion of CEP170 rendered similar resection defect, our data also favors that CEP170 might be the main centrosomal factor accounting for the centrosomes’ role in resection. We then considered that even if CEP170’s role in DDR might require, initially, a centrosomal localization, a possible scenario could be that CEP170 might delocalize to DNA after damage and play a direct role in HR within the nucleus or even at sites of DNA damage. However, an ectopically expressed version of CEP170 tagged with GFP neither relocalized to the nucleus nor was recruited to laser micro-irradiated DNA damage (unpublished observations). Furthermore, CEP170 protein distribution nucleus vs. cytoplasm was also not altered upon irradiation (Figure EV3A), supporting an indirect role of CEP170 in HR repair pathway from the centrosome.

**Figure 3:**
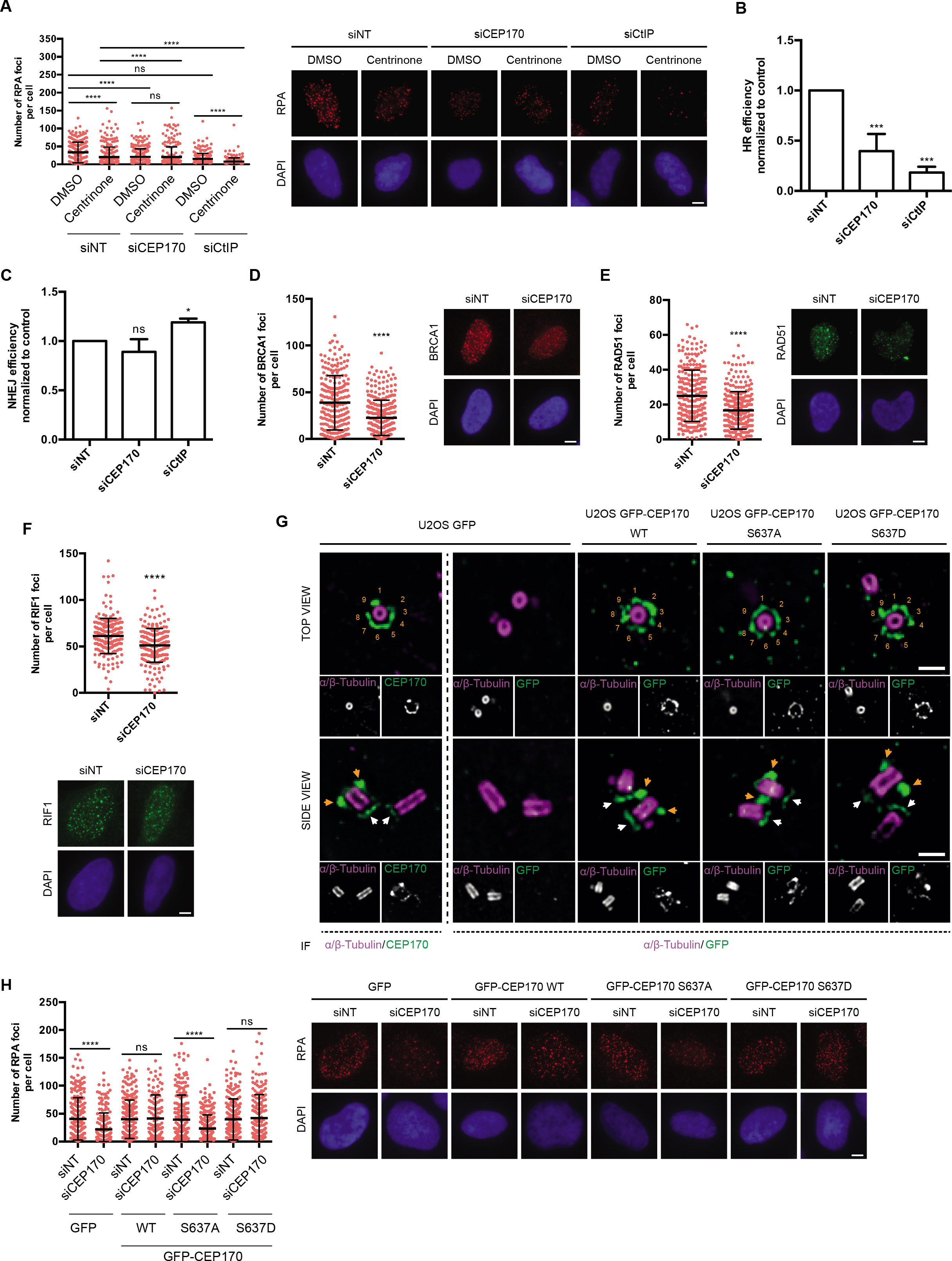
CEP170 phosphorylation at centrosomes is required for resection. **A**, Cells were treated with centrinone or the vehicle DMSO and transfected with the indicated siRNAs. The number of RPA foci per cell for at least 200 cells per condition was quantified automatically using FIJI software and plotted. One representative experiment out of three performed with similar results is shown (left). Representative images are shown (right). Scale bar in white represents 7 μm. Statistical significance was calculated using a two-way ANOVA. **B**, HR was studied as indicated in figure 1A in cells bearing the DR-GFP reporter depleted for the indicated factors. **C**, Same as B, but using cells harboring the NHEJ reporter EJ5-GFP. **D**, The number of BRCA1 foci per cell for at least 200 cells transfected with an siRNA against CEP170 or a siRNA control was quantified automatically using FIJI software and plotted. One representative experiment out of three performed with similar results is shown (left). Representative image on the right side. **E**, Same as D but using an antibody against RAD51 (left). Representative images on the right side. **F**, Same as D but using an antibody against RIF1 (top). Representative images at the bottom. **G**, Ultraextructure expansion microscopy (U-ExM) images of U2OS expressing GFP, GFP-CEP170 WT, GFP-CEP170 S637A or GFP-CEP170 S637D. Images show one representative top view (top panels) and side view (bottom panels). Cells were co-immunostained against alpha and beta tubulin and endogenous CEP170 or GFP as indicated. Orange numbers label the nine SDAs. Orange arrows point to subdistal appendage localization and white arrows to centriole proximal localization of CEP170 variants. Scale bar 500 nm. **H**, same as A but in cells stably expressing the indicated CEP170 variant tagged with GFP or the control GFP plasmid and transfected with siCEP170 or a control sequence (left). Representative images on the right side. In B and C the average and standard deviation of three independent experiments is shown. A-F and H the statistical significance was calculated using a Student’s t-test. p values are represented with one (p < 0.05), two (p < 0.01), three (p < 0.001) or four (p < 0.0001) asterisks. Non statistical significance is labeled ns.

So far, we have shown that the centrosome is required for a proper balance between HR and NHEJ, affecting both repair pathways, and that it specifically regulates the initial step of HR, DNA end resection. Also, that the subdistal centriolar protein CEP170 plays a critical role in DNA processing activation. To study whether the role of CEP170, as this of centrosomes, extend not only to DNA end resection but also to both HR and NHEJ, we analyzed the ability of CEP170 depleted cells to perform different types of DSB repair using the GFP-based reporters described above. In agreement with a role in DNA end resection, homologous recombination Rad51-dependent gene conversion pathway was compromised (Figure 3B). On the other hand, and in stark contrast with our observation in centrosome-less cells, no significant impact on the NHEJ repair pathway was observed (Figure 3C). In addition to the above-mentioned drop of RPA foci in CEP170 depleted cells after irradiation, we also observed a drop of BRCA1 or Rad51 foci (Figure 3D and 3E) and a small but significant reduction on the recruitment of the NHEJ facilitator RIF1 (Figure 3F) along the same lines to the observed changes upon centrinone treatments. Similar changes were observed in CEP128 KO RPE1 cells (Figure EV3B-C). All together our data supports that CEP170, and the SDA, plays a critical role on the centriole subdistal appendages-based regulation of HR. It also suggests that the increase on NHEJ observed in cells lacking centrosomes must be mediated by an additional, yet undiscovered, centriolar factor.

### CEP170-S637 phosphorylation modulates DNA end resection

We then wondered about the molecular steps that might connect DSBs repair and CEP170’s role in HR. Neither cytoplasmic CEP170 protein levels nor overall CEP170 levels changed after irradiation-induced DNA damage (Figure EV3A and D). Remarkably, CEP170 was previously identified in a large-scale proteomic analysis of proteins phosphorylated in response to DNA damage on a consensus site recognized by ATM and ATR (Matsuoka *et al*, 2007). Specifically, Serine-637 of CEP170 is an evolutionary conserved site in vertebrates (Figure EV3E) identified as a residue phosphorylated after DNA damage (Matsuoka *et al*, 2007). To assess if this putative phosphorylation site is biologically relevant, we generated mutated versions of CEP170 either mimicking (CEP170-S637D) or disrupting this phosphorylation (CEP170-S637A) (Figure EV3F). We confirmed by Ultrastructure Expansion Microscopy (U-ExM) that both mutated versions of CEP170 tagged to GFP, when expressed ectopically, showed a similar localization to the subdistal appendages and the proximal side of the centriole as the one reported for the endogenous protein or the also ectopically expressed wildtype version of GFP-tagged CEP170 (Figure 3G). Only in a minority of cells, but regardless of the version of the protein expressed, the overexpression of the protein rendered the accumulation in additional centrosomal locations (Figure EV3G). Thus, we concluded that this expression of different CEP170 species can be used as a complementation system for the endogenous protein. Interestingly, expression of CEP170-S637D but not CEP170-S637A rescued CEP170-depletion RPA phenotype (Figure 3H), suggesting that not only the presence of the protein, but its phosphorylation, is required to fully activate resection. We hypothesized then that after irradiation, CEP170 is phosphorylated by ATM/ATR in S637 which is critical for promoting HR as the pathway of choice for DSB repair. As expected, the expression of CEP170-S637D alone was unable to rescue the drop of RPA foci formation observed upon chemical inhibition of ATM/ATR (Figure EV3H), in agreement with the fact that CEP170 is just one of many targets involved in homologous recombination that is phosphorylated by ATM/ATR in response to DNA damage. As our results support that CEP170’s role in HR is centrosomal-dependent (Figure 3A), and ATM/ATR have been reported to localize to the centrosome (Mullee & Morrison, 2016), we hypothesized that such phosphorylation happens at that localization. We then wondered whether once phosphorylated, CEP170 function in HR could be centrosome independent. To assess this possibility, we overexpressed the phosphomimic CEP170-S637D mutant in centriole depleted cells. Interestingly, CEP170-S637D did not compensate the drop in HR observed in centriole depleted cells (Figure EV3I). Therefore, we concluded that CEP170 location at the centrosome is most likely required for CEP170 phosphorylation by ATM/ATR and for its subsequent role promoting HR.

### Centrosome/CEP170 DNA damage response contributes to cell survival after DNA double strand break

The above-mentioned results reveal a new function of the centrosome and the centriolar protein CEP170 in DSB repair. To assess whether this function contributes to cell survival and proliferation after DNA damage, we performed clonogenic assays with RPE-1 cells either absent of centrioles or heterozygous by CRISPR for CEP170 (Figure EV2K). In agreement with our hypothesis, both conditions rendered cells hypersensitive to DSB-inducing agents such as ionizing radiation (IR), the radiomimetic agent Neocarzinostatin (NCS), or both topoisomerase poisons Camptothecin (CPT) and Etoposide (VP-16) (Figures 4A-H, Figure EV4). These hypersensitivities were of a similar extent to the ones observed for other canonical regulators of HR repair pathway choice such as BRCA1 (Cruz-García *et al*, 2014). Interestingly, CEP170 knockdown, albeit causing a similar reduction on RPA foci and HR as centrinone treatment, rendered a stronger hyper-sensitivity to DNA damaging agents, probably reflecting the lack of stimulation of NHEJ in that condition when compared with centrinone treatment. Notably, this hypersensitivity to irradiation was also observed in CEP170 siRNA depleted cells and could be rescued by expressing the siRNA resistant version of CEP170 (Figure 4I-J). We concluded that the role of the centrosomes and specifically the centriolar protein CEP170 in HR DNA repair is critical to ensure cell survival and proliferation upon DNA damage.

**Figure 4:**
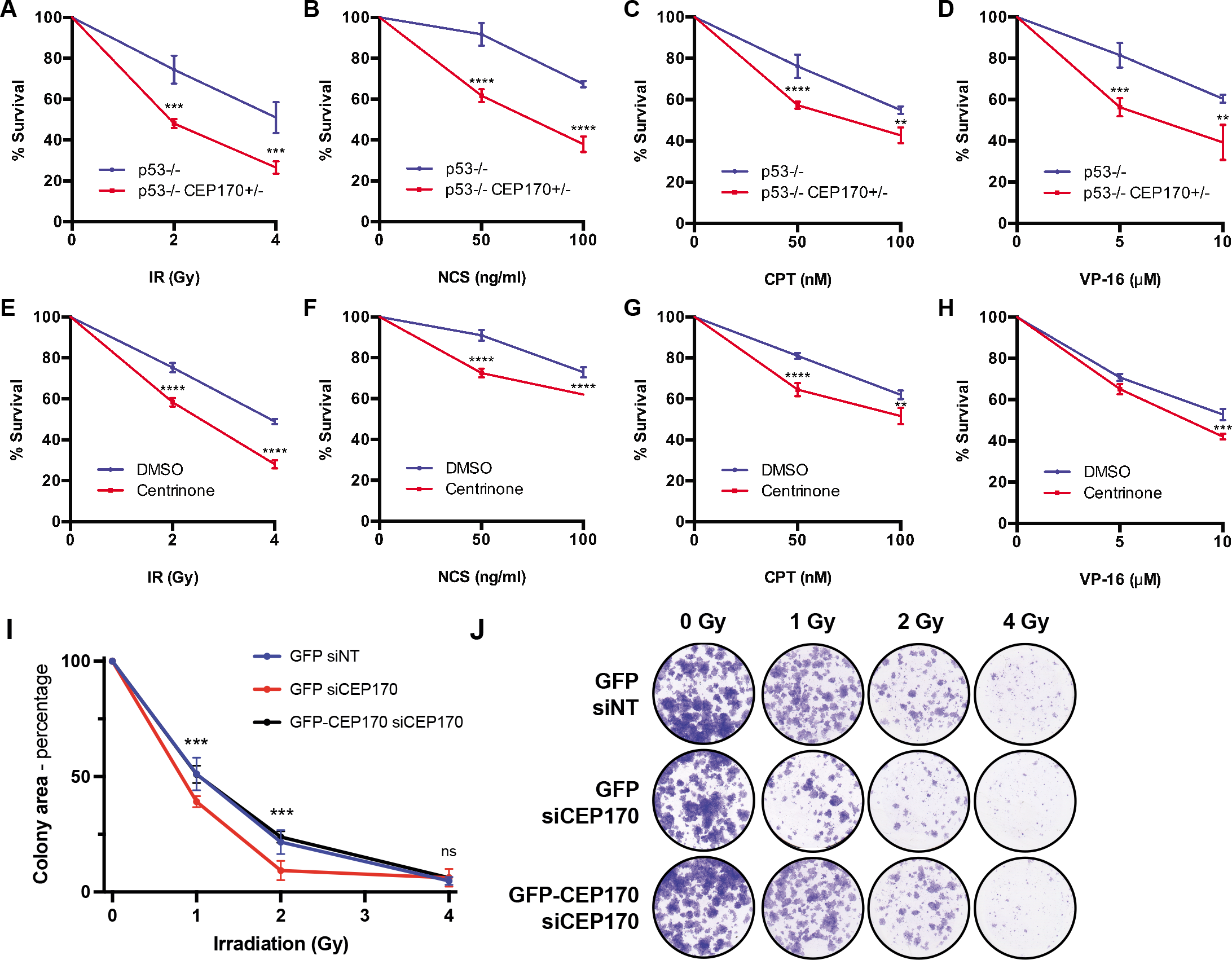
Centrosomes or CEP170 loss hyper-sensitize to treatment with DNA damaging agents. **A**, RPE-1 cells knocked out for p53 and heterozygous for CEP170 (red) or a control RPE-1 p53 KO cells (blue) were exposed to the indicated doses of ionizing radiation. Cell survival was analyzed as described in the methods section using a clonogenic assay. Statistical significance was calculated using a two-way ANOVA. **B**, Same as A but treating cells with the indicated doses of NCS. **C**, Same as A but in cells treated with the indicated doses of CPT. **D**, Same as A but in cells treated with the indicated doses of VP-16. **E**, Same as A but in RPE p53-/ cells treated with centrinone (red) or DMSO (blue) for 7 days prior the treatment with ionizing radiation. **F**, Same as E but in cells treated with the indicated doses of NCS. **G,** Same as E but in cells treated with the indicated doses of CPT. **H**, Same as E but in cells treated with the indicated doses of VP-16. **I**, U2OS cells expressing GFP or GFP-CEP170 and transfected with siRNA against CEP170 or non-target sequence were exposed to ionizing irradiation or mock treated. Cell survival was analyzed as in A. Statistical significance was calculated using a two-way ANOVA. p values are represented with one (p < 0.05), two (p < 0.01), three (p < 0.001) or four (p < 0.0001) asterisks. Non statistical significance is labeled ns. **J**, Representative scanned images of clonogenic assay represented in I. Data shown in all panels are the average of three biological independent experiments each of them performed in three technical replicates.

### CEP170 in cancer biology and cancer prognosis

Strikingly, changes in the overall levels of CEP170, either downregulation or upregulation, can be observed in many tumor samples (Figure 5A). So, we then speculated that, considering this newly found role in controlling DNA repair, the levels of this protein might relate with the appearance of specific mutations during cancer progression. In other words, we wondered whether tumor cells with low levels of CEP170 might contribute to the appearance of specific mutational signatures and, furthermore, whether those signatures might correlate with defects in specific DNA repair pathways, mainly with HR defects. Using the Pan-Cancer Analysis of Whole Genomes (PCAWG) (Campbell *et al*, 2020) we found 435 tumor samples from which we could obtain and analyze mutations on DNA and CEP170 expression data. Based on CEP170 expression, we divided the samples in four quartiles, from the lowest (Q1) to the highest (Q4) RNA levels. Then, we used the SigProfiler software (Bergstrom *et al*, 2019; Islam *et al*, 2020), and identified 16 optimal mutation signatures in our dataset by the trade-off of the mean sample cosine distance and average stability of solutions in the range from 1 to 20 (Figure 5B; Figure EV5A; see methods for details). Out of those 16 signatures, 11(names marked in red in Figure 5C-R) showed significant exposure differences between samples with low CEP170 expression (Q1) versus high levels of the protein (Q4), with a general increase in those specific mutation burdens (Figure 5C-R). The composition of the actual mutational signatures can be found in Figure EV5A. Next, we compared our signatures to the Catalogue of Somatic Mutations in Cancer (COSMIC) (Tate *et al*, 2019) reference mutational signatures to infer the potential sources of the observed mutagenesis (Appendix Figure S1). Strikingly, from this comparison, we observed that from the signatures that showed exposure differences in tumor cells expressing lower levels of CEP170, stood out several that were similar to the ones associated with defective DNA repair in COSMIC, and outstandingly the one related with defective HR, (Figure 5I) in agreement with our observation that this protein is required for proficient recombination. Remarkably, expression levels of other subdistal appendages proteins as NIN and Centriolin showed similar mutational signatures to the one of CEP170, including compatible with defective HR (Figure EV5B and Appendix Figure S2). More interestingly, in this case NDEL1 showed a similar pattern but orders of magnitude less statistically significant, whereas CEP128 did not (see discussion).

**Figure 5:**
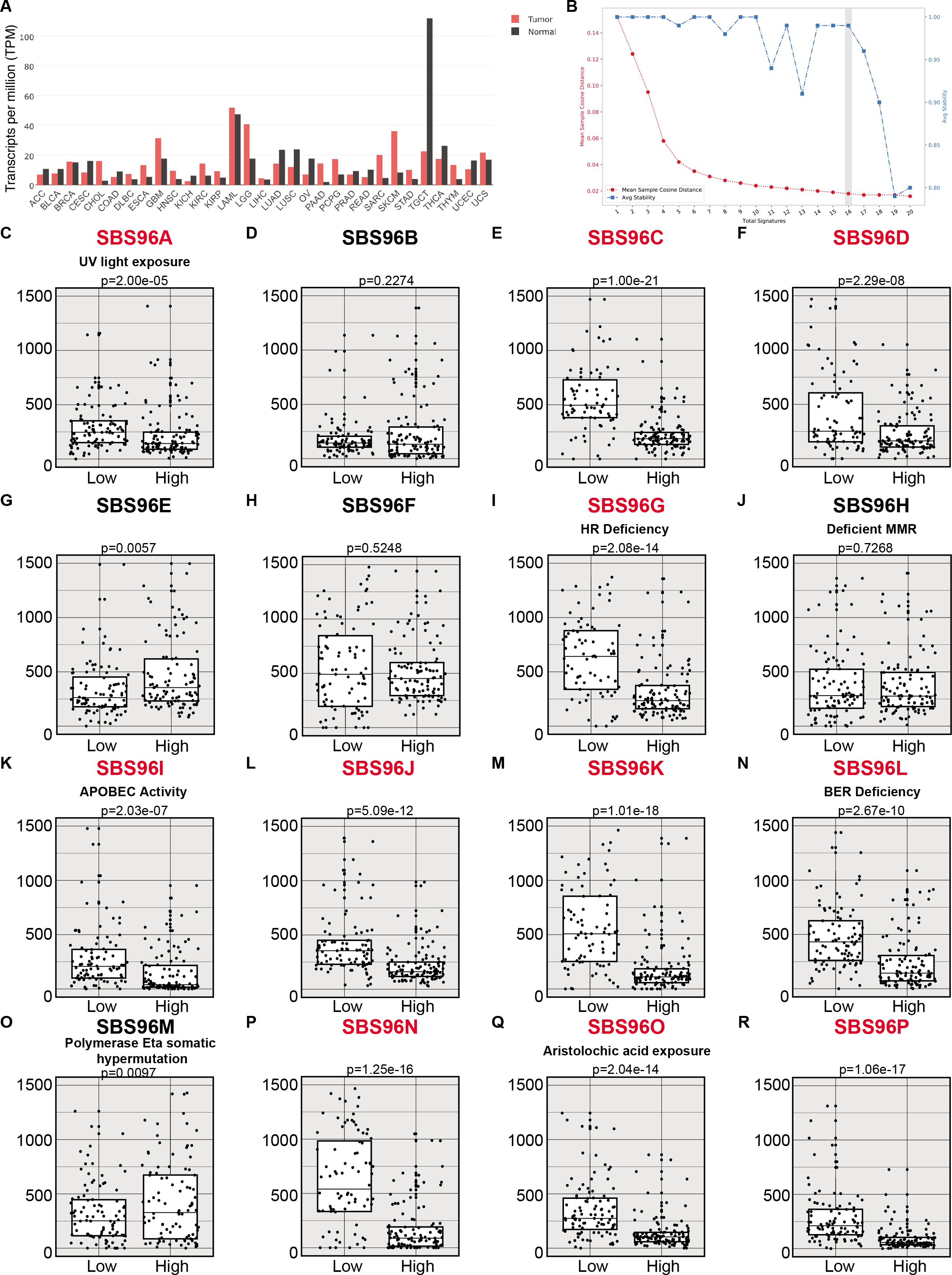
CEP170 levels modulate the mutational signature in cancer samples. **A**, CEP170 levels in tumor samples (red) and the equivalent healthy tissue (black), studied as indicated in the methods section. Tumor acronyms are: ACC Adrenocortical carcinoma; BLCA Bladder urothelial carcinoma; BRCA Breast invasive carcinoma; CESC Cervical squamous cell carcinoma and endocervical adenocarcinoma; CHOL Cholangiocarcinoma; COAD Colon adenocarcinoma; DLBC Lymphoid neoplasm diffuse large B-cell lymphoma; ESCA Esophageal carcinoma; GBM Glioblastoma multiforme; HNSC Head and neck squamous cell carcinoma; KICH Kidney chromophobe; KIRC Kidney renal clear cell carcinoma; KIRP Kidney renal papillary cell carcinoma; LAML Acute myeloid leukemia; LGG Brain lower grade glioma; LHIC Liver hepatocellular carcinoma; LUAD Lung adenocarcinoma; LUSC Lung squamous cell carcinoma; OV Ovarian serous cystadenocarcinoma; PAAD Pancreatic adenocarcinoma; PCPG Pheochromocytoma and Paraganglioma; PRAD Prostate adenocarcinoma; READ Rectum adenocarcinoma; SARC Sarcoma; SKCM Skin cutaneous melanoma; STAD Stomach adenocarcinoma; TGCT Testicular germ cell tumors; THCA Thyroid carcinoma; THYM Thymoma; UCEC Uterine corpus endometrial carcinoma; UCS Uterine carcinosarcoma. **B**, Cancer samples from PCAWG database containing mutation and CEP170 expression data were analyzed using SigProfiler to determine the number of significant signatures that could be analyzed in the dataset. The average stability and Mean Sample Cosine were plotted, and the number of optimal signatures was defined as 16. For other details, see the methods section. **C-R**, Each of the 16 optimal mutational signatures were analyzed in samples at the lower quartile of CEP170 (low) or at the highest (high). The number of mutations in each sample associated to each signature was plotted. The average is shown. Statistical significance was calculated using a Wilcoxon test, and the p-value is shown on top of the graph. The assigned name of the signature is shown on top. The name of those signatures that show statistically significant changes between samples with high and low expression of CEP170 is written in red. For those signatures associated with a known etiology according to the COSMIC database, this is depicted between the name and the p-value.

As cells depleted for CEP170 were hypersensitive to DNA damaging agents used in chemotherapy (Fig 4A-D), and tumor samples with lower levels of CEP170 expression accumulate mutations that are consistent, among other causes, with an HR defect, we hypothesized that the levels of this centrosomal factor might affect the overall survival in cancer patients. In this scenario, low levels of CEP170 could render tumoral cells deficient in HR and, therefore, hypersensitive to anticancer treatments, thus increasing patient survival. Strikingly, we found that this was the case in several tumor types. Indeed, patients with tumors that showed lower levels of CEP170 had a better prognosis when survival was analyzed from patient databases, likely because of a heightened sensitivity to treatment due to an inability to repair proficiently by HR (Figure 6A-L). This suggests that CEP170’s role in HR might be relevant in cancer progression but also its levels might be exploited as a biomarker for cancer treatment response.

**Figure 6:**
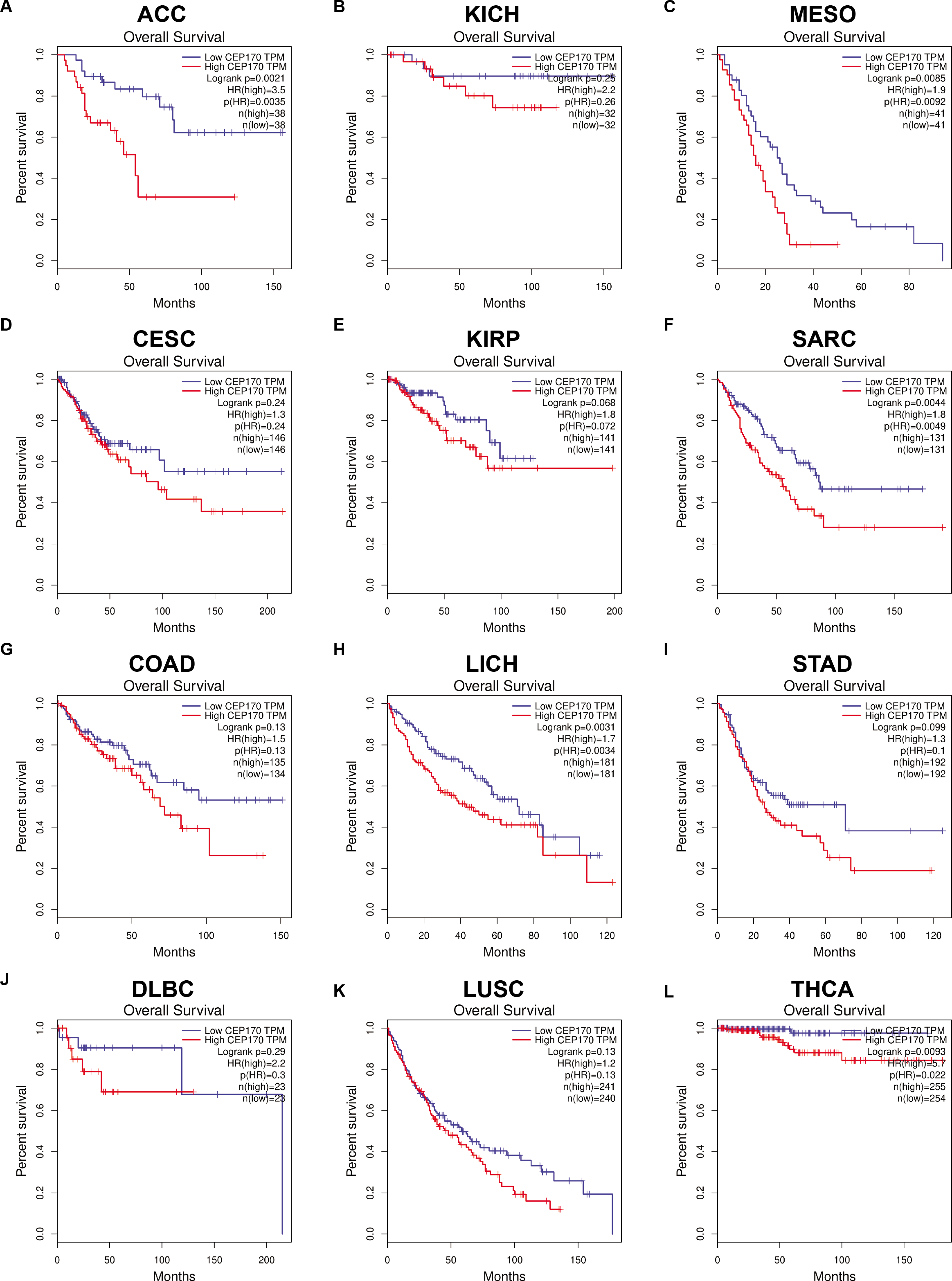
CEP170 levels predict the prognosis of cancer patients. Survival data in different cancer types with different CEP170 expression were collected from the web-based tool GEPIA. Samples were split in two categories, the ones with an expression above the median (high; red lines) or below the median (low; blue lines). Overall survival of each category in each cancer type were plotted in Kaplan-Meyer graphs. Statistical significance was calculated using a Log-rank (Mantel-Cox) test and the p values are included. Tumor types: **A**, ACC Adrenocortical carcinoma. **B**, CESC Cervical squamous cell carcinoma and endocervical adenocarcinoma. **C**, COAD Colon adenocarcinoma. **D**, DLBC Lymphoid neoplasm diffuse large B-cell lymphoma. **E**, KICH Kidney chromophobe. **F**, KIRP Kidney renal papillary cell carcinoma. **G**, LHIC Liver hepatocellular carcinoma. **H**, LUSC Lung squamous cell carcinoma; **I**, MESO Mesothelioma. **J**, SARC Sarcoma. **K**, STAD Stomach adenocarcinoma. **L**, THCA Thyroid carcinoma.

## Discussion

The centrosome is a multifunctional organelle frequently highlighted for its ability to nucleate MTs. Indeed, the more striking consequences of centrioles loss in cell cycle progression is a significant delay in spindle assembly which has been linked to the lack of centrosomal MTOC activity (Wong *et al*, 2015; Fong *et al*, 2016; Meitinger *et al*, 2016; Lambrus *et al*, 2016). Additionally, the centriole works as a scaffold to assemble the primary cilia and flagella, a function evolutionary conserved from the last common eukaryotic ancestor (Bornens, 2012). However, the centriole has also been proposed to play roles unrelated to these activities. For instance, in *C. elegans*, after fertilization, the sperm centrioles provide the initial clue to define the anterior-posterior cell axis and trigger the polarization of the one cell stage embryo required for the first asymmetric embryonic division (Portier *et al*, 2007; Hachet *et al*, 2007). Similar signaling functions of the centrosome has been reported also in mammalian cells (Arquint *et al*, 2014). Several cell cycle regulators localize to the centrosome (Cdk1/Cyclin B, Cdc25B, Cdc25C, Plk1 and Aurora A), which serve as a regulatory platform for G2/M transition, a process that needs to be coordinated to centrosome maturation in order to assemble the mitotic spindle in proper time (Arquint et al., 2014). An additional example on how the centrosome is involved in a signaling pathway to stop cell proliferation of cells with excess number of centrioles has been reported recently. Such role is mediated in human cultured cells by the distal appendage protein ANKRD26, which in turn facilitates the activation of the PIDD as a protective response in cells with excess number of centrioles (Fava *et al*, 2017; Evans *et al*, 2021). Overall, these roles suggest that the centrosome might work as a signaling hub that receives, integrates and elicits a response to preserve cellular viability and fitness.

### Role of the centrosome on DNA damage

It has been previously proposed that the centrosome is important for a fully responsive DNA damage response. This hypothesis is sustained by the fact that several DDR proteins have been found at centrosome and some centrosomal components are directly involved in DDR (Mullee & Morrison, 2016). However, if centriole loss impairs a specific DDR pathway has not been tested. Here, we show that the centrosome directly regulates the fate of DSBs by channeling its repair toward specific pathways. This new role of the centriole affects cells in both S and G2 phases and is independent of centrosomal MTOC activity during interphase. Centrosome loss by itself leads to a bias in DSBs repair towards NHEJ, upregulating this error prone pathways and impairing the more error-free repair by HR. Indeed, overall DSBs repair, measured by γH2AX foci disappearance, was not affected, arguing that the centrosome promotes quality but is not required for the quantity of the repair. Interestingly, such effect does not simply relay on hampering DNA end resection, as the recruitment of the resection antagonist RIF1 protein is not increased, thus suggesting a more active regulatory pathway at play. Despite that, such change in the repair quality is, indeed, relevant for cell fitness, as centriole depleted human cells were hypersensitive to DNA damaging drugs. Remarkably, this hypersensitivity was not observed on genetically modified DT-40 chicken cells depleted of centrioles (Sir *et al*, 2013). The use of drugs targeting centriole duplication has been suggested before for certain types of tumors as breast cancers or gliomas that overexpress TRIM37. Cells from these types of tumors are hypersensitive to centrosome loss (Meitinger *et al*, 2020; Yeow *et al*, 2020). Considering our data, we propose that the same logic could be applied for the treatment of other tumor types where centrosomal loss could be synergistic with DNA damaging drugs for cancer treatment.

### The role of CEP170 in the centrosome-mediated regulation of DSB repair

We have been able to identify a particular substructure within the centriole that plays important roles in the regulation of DNA end resection and recombination: the subdistal appendages. More specifically, the part that branches from the centriole from CEP128 and finishes at CEP170. Furthermore, we found that among these proteins CEP170 appears as a key factor mediating the response, as the degree of resection defect observed upon depletion of subdistal appendage factors directly correlates with the impairment of the recruitment of CEP170. In addition to its subdistal appendage localization CEP170 also localizes to the proximal side of the centriole. Although we cannot rule out that CEP170 might be playing a role in DDR from its centriolar proximal localization, the fact the depletion of factors that specifically interfere with CEP170 recruitment to the subdistal appendages and not to its proximal localization at the centriole (i.e. ODF2 and CEP128 depletion (Hall & Hehnly, 2021)) support the notion that subdistal appendages and more specifically CEP170 from the subdistal appendages are involved in DDR. Subdistal appendages, and specifically CEP170, have previously been involved in MT anchoring to the distal side of the centrioles (Hall & Hehnly, 2021). If this function of CEP170 might be related to its role in DDR is not clear, however we consider this possibility unlikely as drugs impacting MTs dynamics (i.e., Taxol and Nocodazol) did not show DDR defects. Interestingly, a detailed characterization of the behavior of the DSB repair machinery revealed differences between removing the entire centrosome and depleting CEP170. More strikingly, both complete centrosome removal and CEP170 depletion induced a drop of HR, but depletion of CEP170 did not render an increase in NHEJ repair pathway, in stark contrast with cells treated with centrinone. This fact suggests two interesting ideas. First, it reinforces the hypothesis that the upregulation of NHEJ observed upon centrosome loss does not rely on a passive effect due to a resection impairment but rather to an active regulatory process yet to be defined. Therefore, additional centrosomal factors connecting this organelle directly with NHEJ regulation, and independent of the pathway reported in this work, await to be discovered. And second, that CEP170 depletion causes a stronger defect in DSB repair, as the defect in HR is not counterbalanced by an excess in NHEJ, a hypothesis supported by a higher sensitivity to DNA damaging agents. Might other centriolar proteins promote DNA end resection independently of CEP170? We think this is unlikely as treating CEP170 depleted cells with centrinone did not further downregulate HR. Therefore, our data suggest that subdistal appendages through CEP170 is the most relevant pathway that promotes HR for double DNA strand break repair, however still unidentified centrosomal factors might restrict NHEJ repair pathway choice.

### The centrosome, a DDR signaling hub outside the nucleus

But how does the centriole and, specifically CEP170, affect a nuclear event such as DNA end resection without being targeted itself to the nucleus? The easiest explanation would be that the centrosome itself acts as a signaling hub. In this scenario, and similarly to what has been proposed for the full activation of the DDR (Mullee & Morrison, 2016), the DNA damage-elicited signal mediated by ATM/ATR is translated to the centrosome, that will integrate this signal with additional cues from the cellular environment. We propose that a second wave of signaling departs from the centrosome towards the nucleus to regulate DSB repair. Indeed, CEP170 regulatory function on DNA resection readily requires its phosphorylation by ATM/ATR at Ser367, and the phosphorylated protein acts without leaving the centrosome. Furthermore, phosphorylated CEP170 requires the centrosome to elicit its function on DSB repair, supporting a signaling role of this protein from the centrosome. If these phosphorylation affects centriolar functions of CEP170 remains to be tested.

But which factor mediates this second wave? An obvious candidate is BRCA1, which is both a DNA end resection modulator and has been shown to be recruited at the centrosomes. In fact, the sensitivity to DNA damaging agents shown by centrinone treatment or CEP170 depletion is similar to the one observed by the canonical HR regulator BRCA1 (Cruz-García *et al*, 2014). Unfortunately, we have not been able to connect changes in BRCA1 dynamics at the centrosome or in the interaction with CEP170 in response to DNA damage. Thus, the link between the resection machinery and CEP170 remains unknown. How this connection works is still under investigation and will be the subject of further works.

In any case, and based on our results we propose that, as an analogy of how computers work, the centrosome behaves as a cytoplasmic microprocessor we call the <u>C</u>entrosomal <u>P</u>rocessing <u>U</u>nit (CPU). In this model, the centrosome receives inputs from different ongoing cellular processes, including the presence or not of DSBs, integrate them within a single hub and compute them to exert a response that will feed back to modulate the relevant cellular molecular mechanisms, including the DDR and the DNA repair pathways themselves. Among the different functions that have been assigned to the centrosome, the role of this organelle as a signaling platform might be the least characterized of them all. Specially, in mammalian cells, very little examples and almost no molecular mechanism have been reported to date nailing down this centrosome function (Arquint *et al*, 2014). Furthermore, examples reported so far of centrosome signaling function in mammalian cells are restricted to centriole biology related processes (i.e. centriole number control, centrosome maturation or spindle assembly) (Arquint *et al*, 2014; Fava *et al*, 2017; Wong *et al*, 2015). To our knowledge, the data presented here are the first example in mammalian cells of the centrosome itself, and not specific centrosomal factors, acting as a signaling platform in a process totally unrelated to canonical centrosome biology. In the particular case of DSB repair, it is interesting that this process relays, at least in relationship with resection regulation, on subdistal appendages, a centriolar substructure that varies with the cell cycle and that is only present in mature centrioles. This could be potentially used to coordinate the cell cycle stage with the DSB repair pathway of choice.

### Relevance of the CPU role on DNA repair in human health

Fully processive and accurate DNA repair is a critical issue in maintaining a healthy homeostasis at the cellular, tissular and organismal levels. Indeed, and as aforementioned, less than optimal repair is at the root of several hereditary rare diseases but also contributes to cancer development. So, does the role of the centrosome as the CPU, and specifically its function on DNA repair, contribute with the appearance of such diseases? Strikingly, there is one rare disease that clearly connects both the centrosome and the DNA end resection: Seckel Syndrome. For years it has been puzzling that this form of hereditary osteodysplastic dwarfism can be caused either by mutations in the DDR and resection factors (ATR, CtIP, DNA2) (O’Driscoll *et al*, 2004; Qvist *et al*, 2011; Shaheen *et al*, 2014), but also centrosomal proteins (CPAP, CEP63, CEP152, NIN) (Al-Dosari *et al*, 2010; Dauber *et al*, 2012; Sir *et al*, 2011; Kalay *et al*, 2011). Little was understood on how defects in these two apparently unconnected elements were able to cause the same clinical phenotypes, although it has been proposed that this centriolar elements are related with ATR activation. Now, we speculate that is the role of the CPU in DNA end resection might be what actually links the centrosome and Seckel. In this scenario, this syndrome should be considered a bona fide repair-deficient syndrome, what explains why it is clinically similar to other repair-deficient diseases, and that the alteration of the CPU contribution to resection is what causes the syndrome upon mutations of centrosomal proteins. Indeed, this factor can be affecting directly ATR activation at the centrosome but also indirectly by regulating DNA end resection, as this processing creates the ssDNA that is recognized by and activates ATR (Ciccia & Elledge, 2010). This would be a good model to explain that mutations in NIN (Seckel 6 variant), a protein that here we have shown is required for fully processive DNA end resection, are linked to this syndrome.

Moreover, this effect in repair of the CPU should be considered in cancer etiology. Alterations of centrosomal proteins can be found in many cancer types (Gönczy, 2015). The more straightforward way to explain this connection between centrosomal defects and cancer is through chromosome missegregation due to defective spindle organization. Indeed, albeit centrosomes are not completely required for spindle formation, the chances of aberrant mitosis increase when they are not fully functional. Now, we propose that in an additional layer, defective centrosomes not only can affect the appearance of chromosomal instability, but also can affect genetic stability through its role as the CPU in the DNA repair. In agreement, cancer samples with lower CEP170 expression consistently show an accumulation of mutations that belong, among others, to mutational signatures associated with defective DNA repair (BER and HR) or associated with exposure to DNA damaging agents (UV, Aristolochic acid). Intriguingly, expression levels of some but not all centriolar proteins involved in CEP170 recruitment to the subdistal appendages also correlate with this group of molecular signatures (i.e. NIN, Centriolin but not CEP128). Furthermore, expression levels of the subdistal appendages component NDEL1, initially not involved in CEP170 recruitment to the centriole, and which depletion in our hands did not impact HR, also showed a slight but significant correlation with mutational signatures associated to defective DNA repair. Altogether, these data reflect that the role of the centriole in DNA repair might be more complex in the context of cancer, and that compensatory mechanisms might be in place when specific factors such as CEP128 are altered. Overall, our data agrees with the idea that the CPU, and specifically CEP170, acts as a critical hub for regulating DNA repair. Thus, a defective CPU, caused by low levels of CEP170, might contribute to cancer progression increasing the mutation burden of the cells. This opens the door to the possibility that CEP170 acts as a tumor suppressor and its mutations might be considered cancer driver mutations. Our data also suggest that centrosomal defects, including reduced CEP170 levels, hyper-sensitize cells to those treatments *in vitro.* Strikingly, this is also supported by the *in silico* analysis of retrospective studies of cancer samples. Thus, considering collectively that induction of DNA DSBs is a first-line therapeutic approach in cancer and the results exposed in this manuscript, we propose that further understanding of the CPU role in DNA repair will open new research lines and therapeutic avenues in oncology.

## Material and Methods

### Cell lines and growth conditions

U2OS and Saos2 cell lines were grown in DMEM (Sigma-Aldrich) and hTERT-RPE1 cell lines were grown in F12 (Sigma Aldrich). These media were supplemented with 10% fetal bovine serum (Sigma-Aldrich), 2 mM L-glutamine (Sigma-Aldrich), 100 units/ml penicillin, and 100 μg/ml streptomycin (Sigma-Aldrich). For cells expressing GFP or GFP-CEP170 plasmids, standard medium was supplemented with 0.5 mg/ml G418 (Gibco, Invitrogen). hTERT-RPE1 p53-/- cell line was donated by Dr. Rosa Rios’ lab. hTERT-RPE1 p53-/- SAS6-/- was kindly gifted by Dr. Tsou (Wang *et al*, 2015). All cell lines tested negative for mycoplasma contamination. hTERT-RPE1 p53-/- CEP170-/+ cell lines were generated by using CRISPR-Cas9 technology. Single cells were seeded in 96-well plates and left to grow prior to characterization and sequencing. When required, cell viability was determined by performing Trypan Blue (Merck) staining. The percentage of cells negative for Trypan Blue (living cells) was automatically quantified by using DeNovix Cell counter (CellDrop FL-UNLTD).

To obtain centriole-less cells, media was supplemented with 125nM centrinone (Hy-18682, MCE) for at least 7 days. In clonogenic survival experiments, cells were exposed to centrinone for 7 days prior to seeding cells in 6-well plates. Centrinone was present during all the duration of the assay.

For microtubule alteration assays, cells were treated for 1h with 20 μM Taxol and 2h with 1 μg/ml Nocodazol prior to ionizing radiation. DMSO was used as control.

### siRNAs, plasmids and transfections

siRNA duplexes were obtained from Sigma-Aldrich or Dharmacon (Table EV2) and were transfected using RNAiMax Lipofectamine Reagent Mix (Life Technologies), according to the manufacturer’s instructions.

The GFP-CEP170 plasmid (Guarguaglini *et al*, 2005) was obtained from Addgene (#41150). siRNA-resistant and GFP-CEP170 S637A and GFP-CEP170 S637D mutant versions were done using QuikChange Lightning Site-Directed Mutagenesis Kit (#210518). pcDNA5 FRT/TO PLK4-mCherry (Moyer *et al*, 2015) was obtained from Addgene (#80269). Plasmid transfection of U2OS cells was carried out using FuGENE 6 Transfection Reagent (Promega) according to the manufacturer’s protocol.

### HR and NHEJ analysis

U2OS cells bearing a single copy integration of the reporters DR-GFP (Gene conversion) or EJ5-GFP (NHEJ) (Pierce *et al*, 1999; Bennardo *et al*, 2008) were used to analyze the different DSB repair pathways. In all cases, cells were plated in 6-well plates. One day after seeding, cells were transfected with the indicated siRNA and the medium was replaced with a fresh one 24 h later. The next day, each duplicate culture was infected with lentiviral particles containing I-SceI–BFP expression construct at MOI 10 using 8 μg/ml polybrene in 2 ml of DMEM. Then, cells were left to grow for an additional 24 h before changing the medium for fresh DMEM, and 48 h after siRNA transfection cells were washed with PBS, trypsinized, neutralized with DMEM, centrifuged for 3 min at 600 g, fixed with 4% paraformaldehyde for 15 min, and collected by centrifugation. Then, cell pellets were washed once with PBS before resuspension in 150 μl of PBS. Samples were analyzed with a LSRFortessa X-20 (BD) with the BD FACSDiva Software v5.0.3. Four different parameters were considered: side scatter (SSC), forward scatter (FSC), blue fluorescence (407 nm violet laser BP, Filter 450/40), green fluorescence (488 nm blue laser BP Filter 530/30). Finally, the number of green cells from at least 10,000 events positives for blue fluorescence (infected with the I-SceI-BFP construct) was scored. To facilitate the comparison between experiments, this ratio was normalized with siRNA control. At least three completely independent experiments were carried out for each condition and the average and standard deviation are represented.

### Clonogenic cell survival assays

To study cell survival after DNA damage, clonogenic assays were carried out seeding 500 cells in 6-well plates in triplicates. The following day, cells were exposed to DNA damaging agents: DSBs were produced by IR or by acute treatment with topoisomerase inhibitor camptothecin (CPT; Sigma), etoposide (VP16; Sigma) or the radiomimetic agent neocarzinostatin (NCS; Sigma). 2 Gy, 4 Gy or mock treated or incubated for 1 h with 0.01, 0.05, or 0.1 μM CPT, 5, 10 or 20 μM VP-16 or vehicle (DMSO) as control. For NCS, cells were exposed for 1 h to 50 and 100 ng/ml concentrations or water as control. After two washes with PBS, fresh medium was added, and cells were incubated at 37 °C for 7 days to allow colony formation. Afterward, cells were stained and visualized in the solution of 0.5% Crystal Violet (Merck) and 20% ethanol (Merck). Once the colonies were stained, the remaining solution was removed, and plates were washed with water. The surviving percentage at each dose was calculated by dividing the average number of visible colonies in treated versus control (mock-treated or vehicle-treated) dishes. For rescue experiments, U2OS cells were transfected with GFP-CEP170 or GFP plasmid as described. 24h after plasmid transfection, transfected cells were selected using DMEM supplemented with G418 for 7 days. Cells were then transfected again with siRNA against CEP170 or non-target sequences and 500 and 1000 cells were seeded for clonogenic assay in 24-well plates in triplicates. Cells were exposed to ionizing irradiation (1, 2, 4 Gy or mock treated) 48h after transfection and incubated at 37 °C for 10 days to allow cellular growth. The surviving percentage at each condition was then calculated measuring the occupied well area.

In both cases, the experiments were repeated in biological triplicates, and each one in technical replicates.

### SDS-PAGE and western blot analysis

Protein extracts were prepared in 2× Laemmli buffer (4% SDS, 20% glycerol, 125 mM Tris–HCl, pH 6.8) and heated 10 minutes at 100°C. Proteins were resolved by SDS–PAGE and transferred to nitrocellulose membranes (Amersham, MERCK). Membranes were blocked with Odyssey Blocking Buffer (LI-COR) and blotted with the appropriate primary antibody and infrared dyed secondary antibodies (LI-COR) (Tables EV2). Antibodies were prepared in blocking buffer supplemented with 0.1% Tween-20. Membranes were air-dried in the dark and scanned in an Odyssey Infrared Imaging System (LI-COR), and images were analyzed with Image Studio software (LI-COR). Cellular fractionation was performed following literature protocols (Gillotin et al., 2018).

### Immunofluorescence and microscopy

Cells were treated with ionizing radiation or mock treated and incubated. The specific ionizing radiation dose and incubation times are described in each experiment. Then, coverslips were washed once with PBS. The fixation step of the immunofluorescence differs depending on the protein to be visualized. For RPA and BRCA1 foci cells were treated with pre-extraction Buffer (25 mM Tris–HCl, pH 7.5, 50 mM NaCl, 1 mM EDTA, 3 mM MgCl_2_, 300 mM sucrose, and 0.2% Triton X-100) for 5 min on ice and fixed with 4% paraformaldehyde (w/v) in PBS for 15 min. For γH2AX, cells growing on coverslips were treated for 10 min on ice with methanol. For RIF1 and RAD51 foci, cells were fixed with 4% paraformaldehyde (w/v) in PBS for 15 min and treated with 0,2% PBS-Triton X-100. Finally, for centrosomal proteins visualization 10 min fixation with methanol at -20°C was performed. In all cases, after two washes with PBS, cells were blocked for 1 h with 5% FBS in PBS, co-stained with the appropriate primary antibodies (Table EV2) in blocking solution overnight at 4 °C or for 2 h at room temperature, washed again with PBS and then co-immunostained with the appropriate secondary antibodies (Table EV2) in blocking buffer. After washing with PBS and dehydrating the samples with ethanol 100%, coverslips were mounted into glass slides using Vectashield mounting medium with DAPI (Vector Laboratories). Samples were visualized and acquired using Leica AF6000 fluorescence microscope or a Leica DMI8 Thunder fluorescence microscope. The analysis of the number of foci formation in all cases was performed automatically using FIJI software and a Fiji custom macro.

### Ultrastructure expansion microscopy (U-ExM)

Ultrastructure expansion microscopy was performed as described by (Gambarotto *et al*, 2019). Briefly, samples were co-immunostained with antibodies against CEP170 or GFP and a combination of antibodies against alpha and beta tubulin. For imaging, samples were mounted in a poly-D-lysine pre-treated 25 mm coverslip. For image acquisition a Leica DM18 Thunder fluorescence microscope using a 100x oil-immersion objective was used. Images were deconvoluted and brightness and contrast were adjusted to improve data visualization.

### Cell cycle analysis

Cells were fixed with cold 70% ethanol overnight, incubated with 250 μg/ml RNase A (Sigma) and 10 μg/ml propidium iodide (Fluka) at 37 °C for 30 min and analyzed with a LSRFortessa X-20 (BD). Cell cycle distribution data were further analyzed using ModFit LT 5.0 software (Verity Software House Inc).

### Quantitative image-based cytometry (QIBC)

QIBC protocol was adapted from (Michelena & Altmeyer, 2017). Centrinone treated or control cells were grown in DMEM medium supplemented with EdU (10 µM) for 30 minutes and irradiated with 10Gy one hour before fixation. Cells were then washed with PBS, treated with pre-extraction buffer and fixed in A 3.6% paraformaldehyde for 15 minutes at room temperature. Cells were then incubated in 0.5% Triton X-100 in PBS for 15 min, in 2% BSA in PBS for 5 min and in Reaction Cocktail solution (50 mM Tris-HCl, pH 7.5, 150 mM NaCl, 25 mM Cooper reactive, Alexa FluorTM 488 Azide ThermoFisher A10266, and 62.38 mM L-Ascorbic Acid) for 30 minutes. Three consecutive washes with 3% BSA in PBS, wash buffer (0.5 mM EDTA in PBS) and PBS 1× was performed before 1h incubation with blocking buffer (3% BSA, 0,1% Tween in PBS). Samples were immunostained with RPA antibodies as described above. Image acquisition was performed with a Leica DM18 Thunder fluorescence microscope using a 63x oil-immersion objective. A single merged image per condition, containing over 5000 cells, was acquired under non-saturating conditions maintaining the settings across all samples within each biological replica. FiJi software was used to detect nuclei and quantify DAPI and EdU intensity per single cell nucleus and used for the identification of each cell cycle phase. RPA foci number per nucleus was automatically determined using a FiJi custom macro, identifying RPA positive cells as those with more than 15 foci on irradiated cells.

### RNA isolation, reverse transcription and quantitative PCR

RNA extracts were obtained from cells using NZY Total RNA Isolation kit (Nzytech) according to manufacturer’s instructions. RNA concentration was quantified by measuring 260 nm absorbance using a NanoDrop DeNovix DS-11 FX spectrophotometer. To obtain complementary DNA (cDNA), 1 μg RNA was subjected to RQ1 DNase treatment (Promega) prior to reverse transcription reaction using Maxima H Minus First Strand cDNA Synthesis kit (Thermo Scientific) according to manufacturer’s instructions. Quantitative PCR from cDNA was performed to corroborate siRNA-mediated knockdown of several proteins. For this, iTaq Universal SYBR Green Supermix (Bio-Rad) was used following manufacturer’s instructions. DNA primers used for qPCR are listed in Table EV2. qPCR was performed in an Applied Biosystem 7500 FAST Real-Time PCR system. The comparative threshold cycle (Ct) method was used to determine relative transcript levels (Bulletin 5279, Real-Time PCR Applications Guide, Bio-Rad), using β-actin expression as internal control. Expression levels relative to β-actin were determined with the formula 2−ΔΔCt (Livak & Schmittgen, 2001).

### Bioinformatic analysis and data processing

Literature search was performed to obtain different list of genes related to DDR or centriole processes. Both peer-reviewed publications and databases were used (Van Dam *et al*, 2013; Balestra *et al*, 2013; Alves-Cruzeiro *et al*, 2014; Gerdes *et al*, 2009; Kim *et al*, 2010; Li *et al*, 2004; Jakobsen *et al*, 2011; Andersen *et al*, 2003; López-Saavedra *et al*, 2016; Milanowska *et al*, 2011; Arcas *et al*, 2014; Prados-Carvajal *et al*, 2021; Matsuoka *et al*, 2007) as sources for these lists. Knime software was used for the processing and generation of a unique list of DDR-related genes; the same was implemented with centriole-related genes. These two lists were cross checked, generating a list of genes linked to both processes (Table EV1).

Survival plots and gene expression profiles plots in different cancer types with different CEP170 expression were collected from the web-based tool GEPIA (Gene Expression Profiling Interactive Analysis), with open access in: http://gepia.cancer-pku.cn/index.html.

Tumor mutation calls were retrieved from the PCAWG resource, containing 2,658 whole-cancer genomes and their matching normal tissues across 38 tumor types (The ICGC/TCGA Pan-Cancer Analysis of Whole Genomes Consortium, Nature, 2020). To identify mutational signatures, we considered only single nucleotide variants (SNVs). A total of 21628933 SNVs were used to generate a matrix counts of the SBS96 contexts using the SigProfilerMatrixGenerator software(Bergstrom *et al*, 2019). Mutational signatures and its COSMIC reference similarity were inferred via the non-negative matrix factorization algorithm implemented in the SigProfilerExtractor software (Islam *et al*, 2020). Finally, we selected those samples that also contained expression data (435), divided them into quartiles of CEP170 expression and plotted mutational signature activities corresponding to the first quartile (Q1; low) and fourth quartile of expression (Q4; high) as boxplots using Rstudio (RStudio Team, 2022).

### Statistical analysis

Statistical significance was determined with a Student’s t-test or ANOVA as indicated using PRISM software (Graphpad Software, Inc.), except for mutational signature activities comparisons between low and high CEP170 expression were Wilcoxon test was applied using Rstudio (RStudio Team, 2022). Statistically significant differences were labeled with one, two, three or four asterisks for p < 0.05, p < 0.01, p < 0.001 or p < 0.0001, respectively.

### Data Availability

No data related with this project has been deposited in a public database. Original microscopy Images can be found at BioImage Archive

## Supporting information

Experimental View 1

Experimental View 2

Experimental View 3

Experimental View 4

Experimental View 5

## ACKNOWLEDGEMENTS

We thank Maikel Castellano-Pozo and Sonia Jimeno for critical reading of the manuscript. We thank CABIMER microscopy facility and Dr. Paloma Domínguez for her technical support in image acquisition and processing. This work was funded by the R+D+I grant US-1255532 Proyectos I+D+i FEDER Andalucía 2014-2020 from the Junta de Andalucía, the grant P18-RT-1204 from the Consejería de Transformación Económica, Industria, Conocimiento y Universidades, Junta de Andalucía and the grant PID2019-104195G from the Spanish Ministry of Science and Innovation-Agencia Estatal de Investigación/10.13039/501100011033. CABIMER is supported by the regional government of Andalucía (Junta de Andalucía). ADC is funded with FPU fellowships from the Spanish Ministry of Education and RPC is funded with an AECC postdoctoral fellowship. SGP is funded by EMBO Postdoctoral Fellowship ALTF 1039-2021. We also want to thank Paul Guichard and Virginie Hamel (University of Geneva, Department of Cell Biology, Sciences III, Geneva, Switzerland) for their support developing the U-ExM.

## Disclosure and competing interest stamen

The authors declare no actual or perceived conflict of interest.

## Expanded View Figure Legends

**Figure EV 1: Lack of centrioles affect DNA repair by homologous recombination. A-B**, U2OS cells were treated with 125nM of centrinone, or DMSO as a control, for 24 h (A) or 7 days (B). Then, the presence of centrosomes was immunodetected using an antibody against the centriolar proteins γ-tubulin and CP110. The number of cells without centrioles was counted and plotted. **C,** U2OS cells treated for 7 days with centrinone or DMSO, as indicated, were exposed to Camptothecin 1μM for 1 h and prepared for immunofluorescence using an anti-RPA antibody as described in the methods section. The number of RPA foci per cell for at least 200 cells per condition was quantified automatically using FIJI software and plotted. One representative experiment out of three performed with similar results is shown. **D**, RPE-1 cells treated for 7 days with centrinone or DMSO, as indicated, were irradiated with 10 Gy and 1 hour later prepared for immunofluorescence using an anti-RPA antibody as described in the methods section. The number of RPA foci per cell was calculated as in C. **E**, As in D, but in U2OS cells treated for 24h with centrinone or DMSO. **F**, As in D, but in p53 KO or p53 KO HsSAS-6 KO RPE-1 cells **G**, U2OS cells transfected with a plasmid harboring a mCherry-tagged PLK4, or an empty vector as a control, were grown for 3 days. Then, the number of centrosomes was immunodetected using an antibody against the centriolar proteins γ-tubulin and CP110. The number of cells with more than four centrioles was counted and plotted. **H**, Same as Figure 1A, but in cells overexpressing or not PLK4. **I**, Same as Figure 1B, but in cells overexpressing or not PLK4. **J**, U2OS cells treated for 7 days with centrinone or DMSO, as indicated, were prepared for immunofluorescence using an anti-γH2AX antibody as described in the methods section. The number of γH2AX foci per cell for at least 200 cells per condition was quantified automatically using FIJI software and plotted. One representative experiment out of three performed with similar results is shown. **K**, DSB repair kinetic using γH2AX as a proxy. U2OS cells exposed to centrinone or DMSO as a control for 7 days were irradiated (2 Gy). Samples were collected at the indicated time points and γH2AX was immunodetected using a specific antibody as described in the method section. NI indicates a non-irradiated sample taken just before irradiation. The number of γH2AX foci was scored and plotted. **L**, Same as figure 1C, but using an antibody against the NHEJ factor RIF1. **M**, Same as figure 1C, but in cells treated with Taxol, Nocodazol or DMSO. **N**, Cell cycle distribution of samples treated for 7 days with centrinone or DMSO, as indicated. **O**, U2OS cells treated for 7 days with centrinone or DMSO, as indicated, were fixed and the percentage of mitotic cells was quantified based on DAPI staining. At least 400 cells were quantified per condition. **P**, Same as O but cells were immunostained with an antibody against phospho-H3S10. At least 400 cells were scored per condition and the percentage of phospho-H3S10 positive cells was quantified. **Q**, Cell viability of U2OS cells treated for 7 days with centrinone or DMSO was quantified by Trypan Blue staining and plotted. **R-T**, Same as Figure 1C-E but using Saos-2 cells. **U-V**, Same as Figure 1C-E but using RPE-1 p53 KO cells. **X-Z**, Same as Figure 1C-E but using RPE-1 p21 KO cells. **A, B, G-I, K, O-Q**, the average and standard deviation of three independent experiments is shown. The statistical significance was calculated using a Student’s t-test. p values are represented with one (p < 0.05), two (p < 0.01), three (p < 0.001) or four (p < 0.0001) asterisks. Non statistical significance is labeled ns.

**Figure EV 2: Identification of the subdistal appendages as involved in HR regulation. A**, Pipeline for identification of candidate genes involved in the connection between the centrosome and the DDR. **B**, U2OS cells were transfected with the indicated siRNAs and 48 h later, protein samples were prepared, resolved in SDS-PAGE and blotted with the indicated antibodies. **C**, U2OS cells were transfected with the indicated siRNAs. 48 h later, total RNA was isolated and the levels of the Ndel1 RNA was calculated using Q-PCR. The levels were normalized to the sample transfected with a control siRNA, taken as one. The average and standard deviation of three experiments is plotted. Statistical significance was calculated using a Student’s t-test. **D**, Same as C but in cells transfected with an siRNA against CEP128 and using Q-PCR primers against the mRNA for this protein. **E**, same as Figure 2D but for RPE-1 CEP128 KO or RPE-1 control cells. **F**, same as Figure 1C but for RPE-1 CEP128 KO or RPE-1 control cells. **G**, protein samples were prepared from cells treated as in Figure 2F, resolved in SDS-PAGE and blotted with the indicated antibodies. **H**, same as Figure EV1C but in CEP170 siRNA depleted or siRNA control cells. **I and J**, same as in Figure EV1J but for CEP170 siRNA depleted or siRNA control cells stained with anti-RPA antibodies (I) or γH2AX antibodies (J). **K**, protein samples were prepared from U2OS CEP170 heterozygous KO clones, resolved in SDS-PAGE and blotted with the indicated antibodies. **C and D**, the average and standard deviation of three independent experiments is shown. The statistical significance was calculated using a Student’s t-test. p values are represented with one (p < 0.05), two (p < 0.01), three (p < 0.001) or four (p < 0.0001) asterisks. Non statistical significance is labeled ns.

**Figure EV 3: CEP170 in response to DNA damage. A**, U2OS cells stably expressing a GFP-CEP170 construct were irradiated (10 Gy, plus sign) or not (minus sign). Protein samples were prepared 1 h later by nuclear fractionation as described in the methods section. Samples were resolved in SDS-PAGE and blotted with the indicated antibodies. Size ladder marker is shown on the right side. A representative image of three independent experiments is shown. **B and C**, same as Figure 1 D and E but for RPE-1 CEP128 KO or RPE-1 control cells. **D**, Total levels of endogenous CEP170 were immunodetected in protein samples from U2OS cells irradiated (10Gy, +IR) or not (-IR) resolved in SDS-PAGE using an antibody against this protein. Size ladder marker is shown on the right side. A representative image of four independent experiments is shown on the top, and the average and standard deviation of the quantification of the western blots at the bottom. **E**, alignment of human CEP170 sequence flanking Serine-637 (red box) with the homologous sequences from the indicated vertebrates. **F**, protein samples were prepared from cells treated as in Figure 3H, resolved in SDS-PAGE and blotted with the indicated antibodies. **G**, Same as in Figure 3G but examples of ectopic centrosomal localization of GFP tagged CEP170 WT and mutant versions. **H**, Same as Figure 3H, but in cells pretreated for 1 hour with inhibitors against ATM (ATMi, white bars), ATR (ATRi, grey bars) or DMSO as a control (left). Representative images are shown on the right. **I**, Same as Figure 1C but in cells treated with centrinone or DMSO and stably expressing the indicated GFP constructs. The statistical significance was calculated using a Student’s t-test. p values are represented with one (p < 0.05), two (p < 0.01), three (p < 0.001) or four (p < 0.0001) asterisks. Non statistical significance is labeled ns.

**Figure EV 4: Centrosomes or CEP170 loss hyper-sensitize to treatment with DNA damaging agents.**

Representative images of clonogenic assays shown in Figure 4A-H

**Figure EV 5: Mutational signatures analyzed in tumor samples with low or high expression of CEP170. A**, for each signature analyzed, the distribution of single base substitution considering the previous and following base is shown. Each signature presents a unique profile of mutations. **B**, same as in Figure 5I but for samples at the lower quartile (low) or at the highest (high) of the indicated centriolar subdistal appendages proteins.

## References

Al-Dosari MS, Shaheen R, Colak D & Alkuraya FS (2010) Novel CENPJ mutation causes Seckel syndrome. J Med Genet 47: 411–414

Alderton GK, Galbiati L, Griffith E, Surinya KH, Neitzel H, Jackson AP, Jeggo PA & O’Driscoll M (2006) Regulation of mitotic entry by microcephalin and its overlap with ATR signalling. Nat Cell Biol 8: 725–733

Alves-Cruzeiro JMDC, Nogales-Cadenas R & Pascual-Montano AD (2014) CentrosomeDB: A new generation of the centrosomal proteins database for Human and Drosophila melanogaster. Nucleic Acids Res 42: 430–436

Andersen JS, Wilkinson CJ, Mayor T, Mortensen P, Nigg EA & Mann M (2003) Proteomic characterization of the human centrosome by protein correlation profiling. Nature 426: 570–574

Arcas A, Fernández-Capetillo O, Cases I & Rojas AM (2014) Emergence and evolutionary analysis of the human DDR network: Implications in comparative genomics and downstream analyses. Mol Biol Evol 31: 940–961

Arquint C, Gabryjonczyk AM & Nigg EA (2014) Centrosomes as signalling centres. Philos Trans R Soc B Biol Sci 369

Balestra FR, Strnad P, Flückiger I & Gönczy P (2013) Discovering regulators of centriole biogenesis through siRNA-based functional genomics in human cells. Dev Cell 25: 555–571

Bennardo N, Cheng A, Huang N & Stark JM (2008) Alternative-NHEJ is a mechanistically distinct pathway of mammalian chromosome break repair. PLoS Genet 4

Bergstrom EN, Huang MN, Mahto U, Barnes M, Stratton MR, Rozen SG & Alexandrov LB (2019) SigProfilerMatrixGenerator: A tool for visualizing and exploring patterns of small mutational events. BMC Genomics 20: 1–12

Blackford AN & Jackson SP (2017) ATM, ATR, and DNA-PK: The Trinity at the Heart of the DNA Damage Response. Mol Cell 66: 801–817

Bornens M (2012) The centrosome in cells and organisms. Science (80-) 335: 422–426

Campbell PJ, Getz G, Korbel JO, Stuart JM, Jennings JL, Stein LD, Perry MD, Nahal-Bose HK, Ouellette BFF, Li CH, et al (2020) Pan-cancer analysis of whole genomes. Nature 578: 82–93

Cejka P (2015) DNA end resection: Nucleases team up with the right partners to initiate homologous recombination. J Biol Chem 290: 22931–22938

Chang HHY, Pannunzio NR, Adachi N & Lieber MR (2017) Non-homologous DNA end joining and alternative pathways to double-strand break repair. Nat Rev Mol Cell Biol 18: 495–506

Ciccia A & Elledge SJ (2010) The DNA Damage Response: Making It Safe to Play with Knives. Mol Cell 40: 179–204

Cruz-García A, López-Saavedra A & Huertas P (2014) BRCA1 accelerates CtIP-ediated DNA-end resection. Cell Rep 9: 451–459

Daly OM, Gaboriau D, Karakaya K, King S, Dantas TJ, Lalor P, Dockery P, Krämer A & Morrison CG (2016) CEP164-null cells generated by genome editing show a ciliation defect with intact DNA repair capacity. J Cell Sci 129: 1769–1774

Van Dam TJP, Wheway G, Slaats GG, Huynen MA & Giles RH (2013) The SYSCILIA gold standard (SCGSv1) of known ciliary components and its applications within a systems biology consortium. Cilia 2: 1–5

Dantas TJ, Wang Y, Lalor P, Dockery P & Morrison CG (2011) Defective nucleotide excision repair with normal centrosome structures and functions in the absence of all vertebrate centrins. J Cell Biol 193: 307–318

Dauber A, Lafranchi SH, Maliga Z, Lui JC, Moon JE, McDeed C, Henke K, Zonana J, Kingman GA, Pers TH, et al (2012) Novel microcephalic primordial dwarfism disorder associated with variants in the centrosomal protein ninein. J Clin Endocrinol Metab 97: E2140–51

Evans LT, Anglen T, Scott P, Lukasik K, Loncarek J & Holland AJ (2021) ANKRD26 recruits PIDD1 to centriolar distal appendages to activate the PIDDosome following centrosome amplification. EMBO J 40: 1–18

Fava LL, Schuler F, Sladky V, Haschka MD, Soratroi C, Eiterer L, Demetz E, Weiss G, Geley S, Nigg EA, et al (2017) The PIDDosome activates p53 in response to supernumerary centrosomes. Genes Dev 31: 34–45

Fong CS, Mazo G, Das T, Goodman J, Kim M, O’Rourke BP, Izquierdo D & Tsou MFB (2016) 53BP1 and USP28 mediate p53-dependent cell cycle arrest in response to centrosome loss and prolonged mitosis. Elife 5: 1–18

Gambarotto D, Zwettler FU, Le Guennec M, Schmidt-Cernohorska M, Fortun D, Borgers S, Heine J, Schloetel JG, Reuss M, Unser M, et al (2019) Imaging cellular ultrastructures using expansion microscopy (U-ExM). Nat Methods 16: 71–74

Gerdes JM, Davis EE & Katsanis N (2009) The Vertebrate Primary Cilium in Development, Homeostasis, and Disease. Cell 137: 32–45

Gönczy P (2015) Centrosomes and cancer: Revisiting a long-standing relationship. Nat Rev Cancer 15: 639–652

Gönczy P & Hatzopoulos GN (2019) Centriole assembly at a glance. J Cell Sci 132

Griffith E, Walker S, Martin CA, Vagnarelli P, Stiff T, Vernay B, Sanna N Al, Saggar A, Hamel B, Earnshaw WC, et al (2008) Mutations in pericentrin cause Seckel syndrome with defective ATR-dependent DNA damage signaling. Nat Genet 40: 232–236

Guarguaglini G, Duncan PI, Stierhof YD, Holmström T, Duensing S & Nigg EA (2005) The forkhead-associated domain protein Cep170 interacts with Polo-like kinase 1 and serves as a marker for mature centrioles. Mol Biol Cell 16: 1095– 107

Le Guennec M, Klena N, Aeschlimann G, Hamel V & Guichard P (2021) Overview of the centriole architecture. Curr Opin Struct Biol 66: 58–65

Hachet V, Canard C & Gönczy P (2007) Centrosomes Promote Timely Mitotic Entry in C. elegans Embryos. Dev Cell 12: 531–541

Hall NA & Hehnly H (2021) A centriole’s subdistal appendages: contributions to cell division, ciliogenesis and differentiation. Open Biol 11: 200399

Hauge S, Macurek L & Syljuåsen RG (2019) p21 limits S phase DNA damage caused by the Wee1 inhibitor MK1775. Cell Cycle 18: 834–847

Huertas P (2010) DNA resection in eukaryotes: Deciding how to fix the break. Nat Struct Mol Biol 17: 11–16

Islam SMA, Wu Y, Díaz-Gay M, Bergstrom EN, He Y, Barnes M, Vella M, Wang J, Teague JW, Clapham P, et al (2020) Uncovering novel mutational signatures by de novo extraction with SigProfilerExtractor. bioRxiv

Jackson SP & Bartek J (2009) The DNA-damage response in human biology and disease. Nature 461: 1071–1078

Jakobsen L, Vanselow K, Skogs M, Toyoda Y, Lundberg E, Poser I, Falkenby LG, Bennetzen M, Westendorf J, Nigg EA, et al (2011) Novel asymmetrically localizing components of human centrosomes identified by complementary proteomics methods. EMBO J 30: 1520–1535

Jasin M & Rothstein R (2013) Repair of strand breaks by homologous recombination. Cold Spring Harb Perspect Biol 5

Kalay E, Yigit G, Aslan Y, Brown KE, Pohl E, Bicknell LS, Kayserili H, Li Y, Tüysüz B, Nürnberg G, et al (2011) CEP152 is a genome maintenance protein disrupted in Seckel syndrome. Nat Genet 43: 23–26

Kim J, Lee JE, Heynen-Genel S, Suyama E, Ono K, Lee K, Ideker T, Aza-Blanc P & Gleeson JG (2010) Functional genomic screen for modulators of ciliogenesis and cilium length. Nature 464: 1048–1051

Lambrus BG, Daggubati V, Uetake Y, Scott PM, Clutario KM, Sluder G & Holland AJ (2016) A USP28-53BP1-p53-p21 signaling axis arrests growth after centrosome loss or prolonged mitosis. J Cell Biol 214: 143–153

Lambrus BG, Uetake Y, Clutario KM, Daggubati V, Snyder M, Sluder G & Holland AJ (2015) P53 protects against genome instability following centriole duplication failure. J Cell Biol 210: 63–77

Leidel S, Delattre M, Cerutti L, Baumer K & Gönczy P (2005) SAS-6 defines a protein family required for centrosome duplication in C. elegans and in human cells. Nat Cell Biol 7: 115–125

Li JB, Gerdes JM, Haycraft CJ, Fan Y, Teslovich TM, May-Simera H, Li H, Blacque OE, Li L, Leitch CC, et al (2004) Comparative genomics identifies a flagellar and basal body proteome that includes the BBS5 human disease gene. Cell 117: 541–552

Livak KJ & Schmittgen TD (2001) Analysis of relative gene expression data using real-time quantitative PCR and the 2-ΔΔCT method. Methods 25: 402–408

López-Saavedra A, Gómez-Cabello D, Domínguez-Sánchez MS, Mejías-Navarro F, Fernández-Ávila MJ, Dinant C, Martínez-Macías MI, Bartek J & Huertas P (2016) A genome-wide screening uncovers the role of CCAR2 as an antagonist of DNA end resection. Nat Commun 7

Matsuoka S, Ballif BA, Smogorzewska A, McDonald ER, Hurov KE, Luo J, Bakalarski CE, Zhao Z, Solimini N, Lerenthal Y, et al (2007) ATM and ATR substrate analysis reveals extensive protein networks responsive to DNA damage. Science (80-) 316: 1160–1166

Meitinger F, Anzola J V., Kaulich M, Richardson A, Stender JD, Benner C, Glass CK, Dowdy SF, Desai A, Shiau AK, et al (2016) 53BP1 and USP28 mediate p53 activation and G1 arrest after centrosome loss or extended mitotic duration. J Cell Biol 214: 155–166

Meitinger F, Ohta M, Lee KY, Watanabe S, Davis RL, Anzola J V., Kabeche R, Jenkins DA, Shiau AK, Desai A, et al (2020) TRIM37 controls cancer-specific vulnerability to PLK4 inhibition. Nature 585: 440–446

Michelena J & Altmeyer M (2017) Cell Cycle Resolved Measurements of Poly(ADP-Ribose) Formation and DNA Damage Signaling by Quantitative Image-Based Cytometry. In Methods in Molecular Biology pp 57–68.

Milanowska K, Krwawicz J, Papaj G, Kosiński J, Poleszak K, Lesiak J, Osińska E, Rother K & Bujnicki JM (2011) REPAIRtoire-A database of DNA repair pathways. Nucleic Acids Res 39: 788–792

Mönnich M, Borgeskov L, Breslin L, Jakobsen L, Rogowski M, Doganli C, Schrøder JM, Mogensen JB, Blinkenkjær L, Harder LM, et al (2018) CEP128 Localizes to the Subdistal Appendages of the Mother Centriole and Regulates TGF-β/BMP Signaling at the Primary Cilium. Cell Rep 22: 2584–2592

Moyer TC, Clutario KM, Lambrus BG, Daggubati V & Holland AJ (2015) Binding of STIL to Plk4 activates kinase activity to promote centriole assembly. J Cell Biol 209: 863–878

Mullee LI & Morrison CG (2016) Centrosomes in the DNA damage response—the hub outside the centre. Chromosom Res 24: 35–51

Nabais C, Peneda C & Bettencourt-Dias M (2020) Evolution of centriole assembly. Curr Biol 30: R494–R502

Nigg EA & Raff JW (2009) Centrioles, Centrosomes, and Cilia in Health and Disease. Cell 139: 663–678

O’Driscoll M, Gennery AR, Seidel J, Concannon P & Jeggo PA (2004) An overview of three new disorders associated with genetic instability: LIG4 syndrome, RS-SCID and ATR-Seckel syndrome. DNA Repair (Amst) 3: 1227–1235

Pierce AJ, Johnson RD, Thompson LH & Jasin M (1999) XRCC3 promotes homology-directed repair of DNA damage in mammalian cells service XRCC3 promotes homology-directed repair of DNA damage in mammalian cells. 2633–2638

Portier N, Audhya A, Maddox PS, Green RA, Dammermann A, Desai A & Oegema K (2007) A Microtubule-Independent Role for Centrosomes and Aurora A in Nuclear Envelope Breakdown. Dev Cell 12: 515–529

Prados-Carvajal R, Rodríguez-Real G, Gutierrez-Pozo G & Huertas P (2021) CtIP-mediated alternative mRNA splicing fine-tunes the DNA damage response. Rna 27: 303–323

Qvist P, Huertas P, Jimeno S, Nyegaard M, Hassan MJ, Jackson SP & Borglum AD (2011) CtIP Mutations Cause Seckel and Jawad Syndromes. PLoS Genet 7: e1002310

Ranjha L, Howard SM & Cejka P (2018) Main steps in DNA double-strand break repair: an introduction to homologous recombination and related processes. Chromosoma 127: 187–214

Shaheen R, Faqeih E, Ansari S, Abdel-Salam G, Al-Hassnan ZN, Al-Shidi T, Alomar R, Sogaty S & Alkuraya FS (2014) Genomic analysis of primordial dwarfism reveals novel disease genes. Genome Res 24: 291–299

Sir J-H, Barr AR, Nicholas AK, Carvalho OP, Khurshid M, Sossick A, Reichelt S, D’Santos C, Woods CG & Gergely F (2011) A primary microcephaly protein complex forms a ring around parental centrioles. Nat Genet 43: 1147–1153

Sir JH, Pütz M, Daly O, Morrison CG, Dunning M, Kilmartin J V. & Gergely F (2013) Loss of centrioles causes chromosomal instability in vertebrate somatic cells. J Cell Biol 203: 747–756

Sivasubramaniam S, Sun X, Pan YR, Wang S & Lee EYHP (2008) Cep164 is a mediator protein required for the maintenance of genomic stability through modulation of MDC1, RPA, and CHK1. Genes Dev 22: 587–600

Symington LS (2016) Mechanism and regulation of DNA end resection in eukaryotes. Crit Rev Biochem Mol Biol 51: 195–212

Tate JG, Bamford S, Jubb HC, Sondka Z, Beare DM, Bindal N, Boutselakis H, Cole CG, Creatore C, Dawson E, et al (2019) COSMIC: The Catalogue Of Somatic Mutations In Cancer. Nucleic Acids Res 47: D941–D947

Tubbs A & Nussenzweig A (2017) Endogenous DNA Damage as a Source of Genomic Instability in Cancer. Cell 168: 644–656

Wang WJ, Acehan D, Kao CH, Jane WN, Uryu K & Tsou MFB (2015) De novo centriole formation in human cells is error-prone and does not require SAS-6 self-assembly. Elife 4: 1–13

Wong YL, Anzola J V., Davis RL, Yoon M, Motamedi A, Kroll A, Seo CP, Hsia JE, Kim SK, Mitchell JW, et al (2015) Reversible centriole depletion with an inhibitor of Polo-like kinase 4. Science (80-) 348: 1155–1160

Yeow ZY, Lambrus BG, Marlow R, Zhan KH, Durin MA, Evans LT, Scott PM, Phan T, Park E, Ruiz LA, et al (2020) Targeting TRIM37-driven centrosome dysfunction in 17q23-amplified breast cancer. Nature 585: 447–452

