## Supplementary figures and images for "Centriolar subdistal appendages promote double strand break repair through homologous recombination"

### Experimental View 1

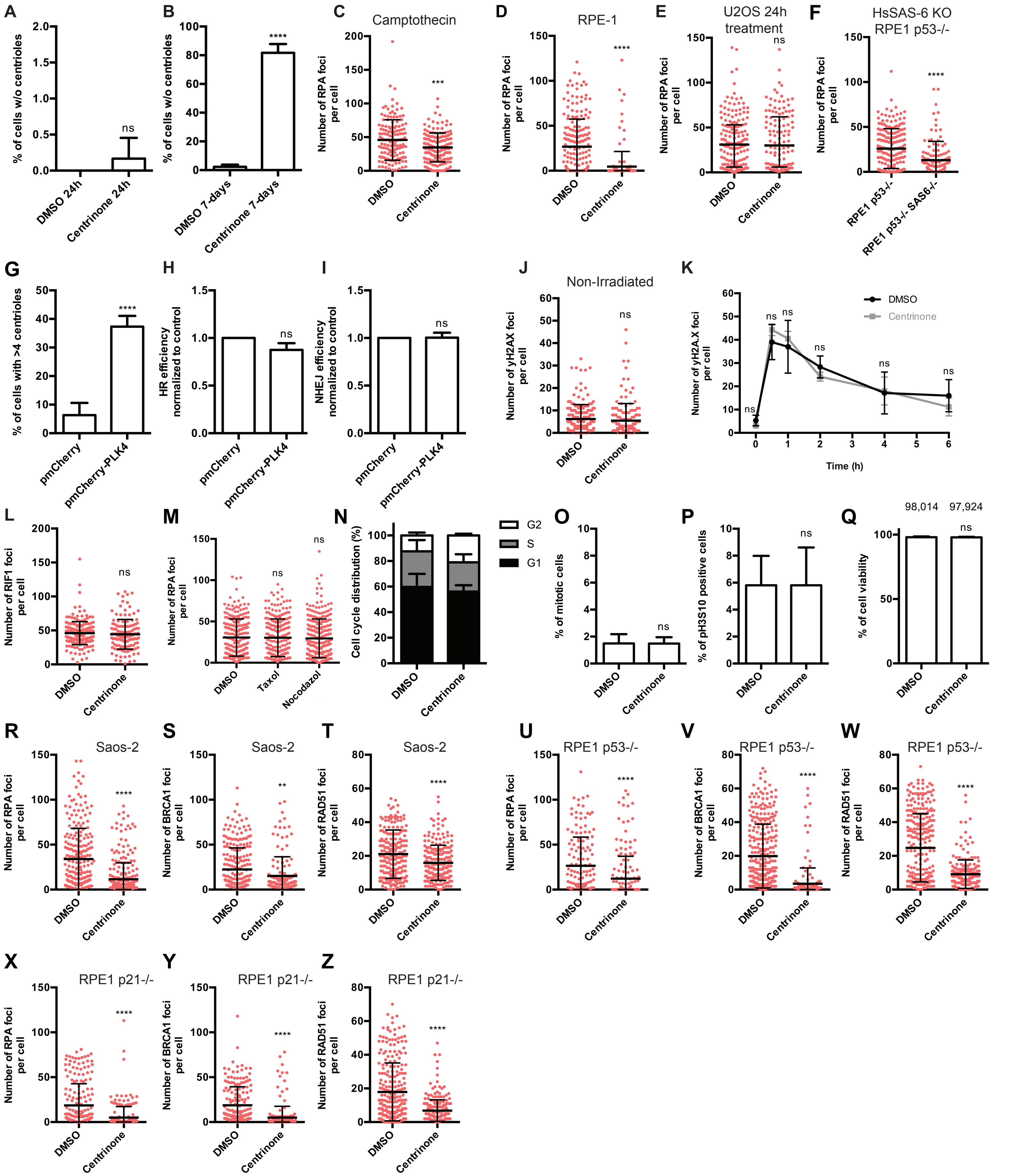


**Rodriguez-Real Figure Expanded View 1**

### Experimental View 2

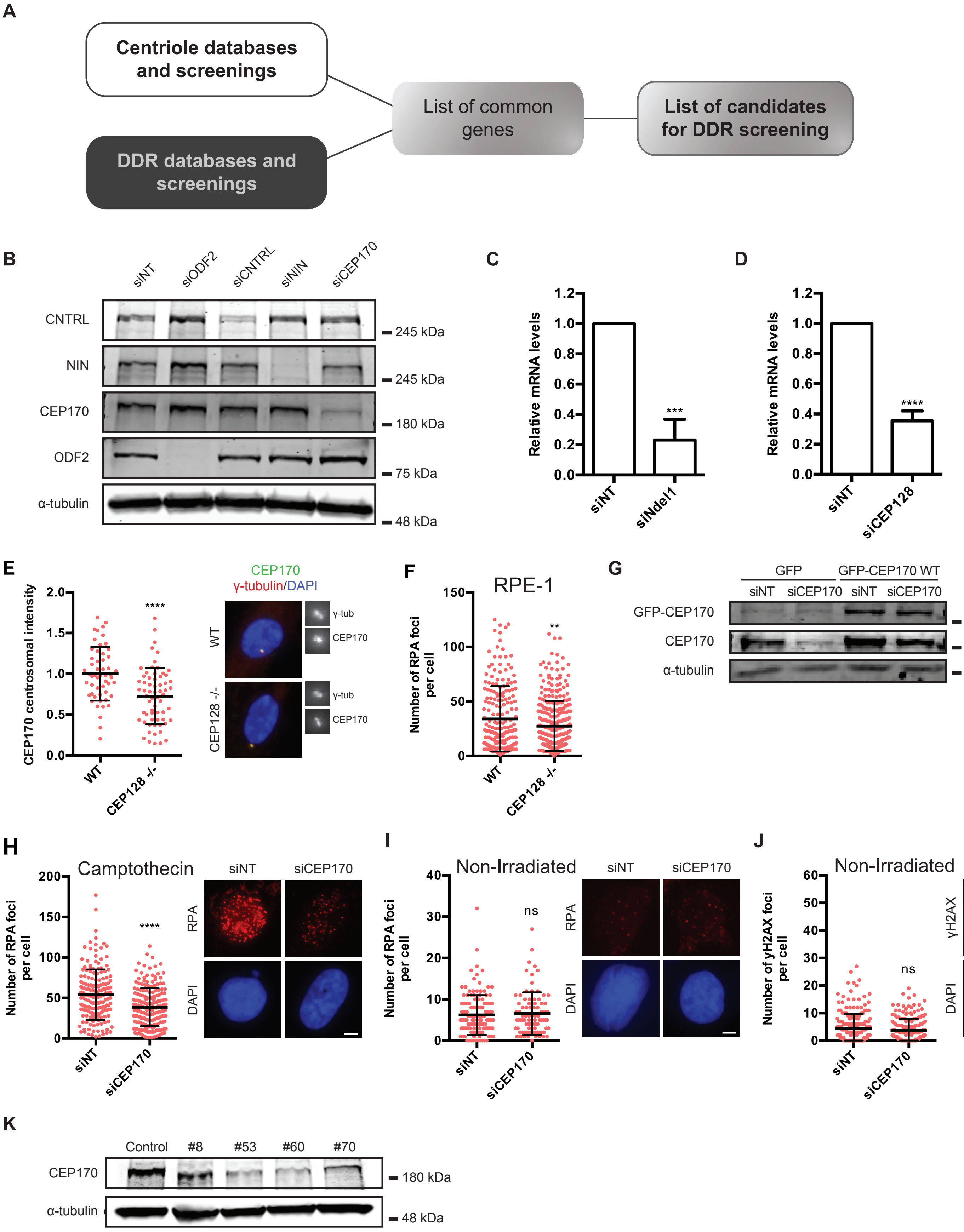


Rodriguez-Real Figure Expanded View 2

### Experimental View 3

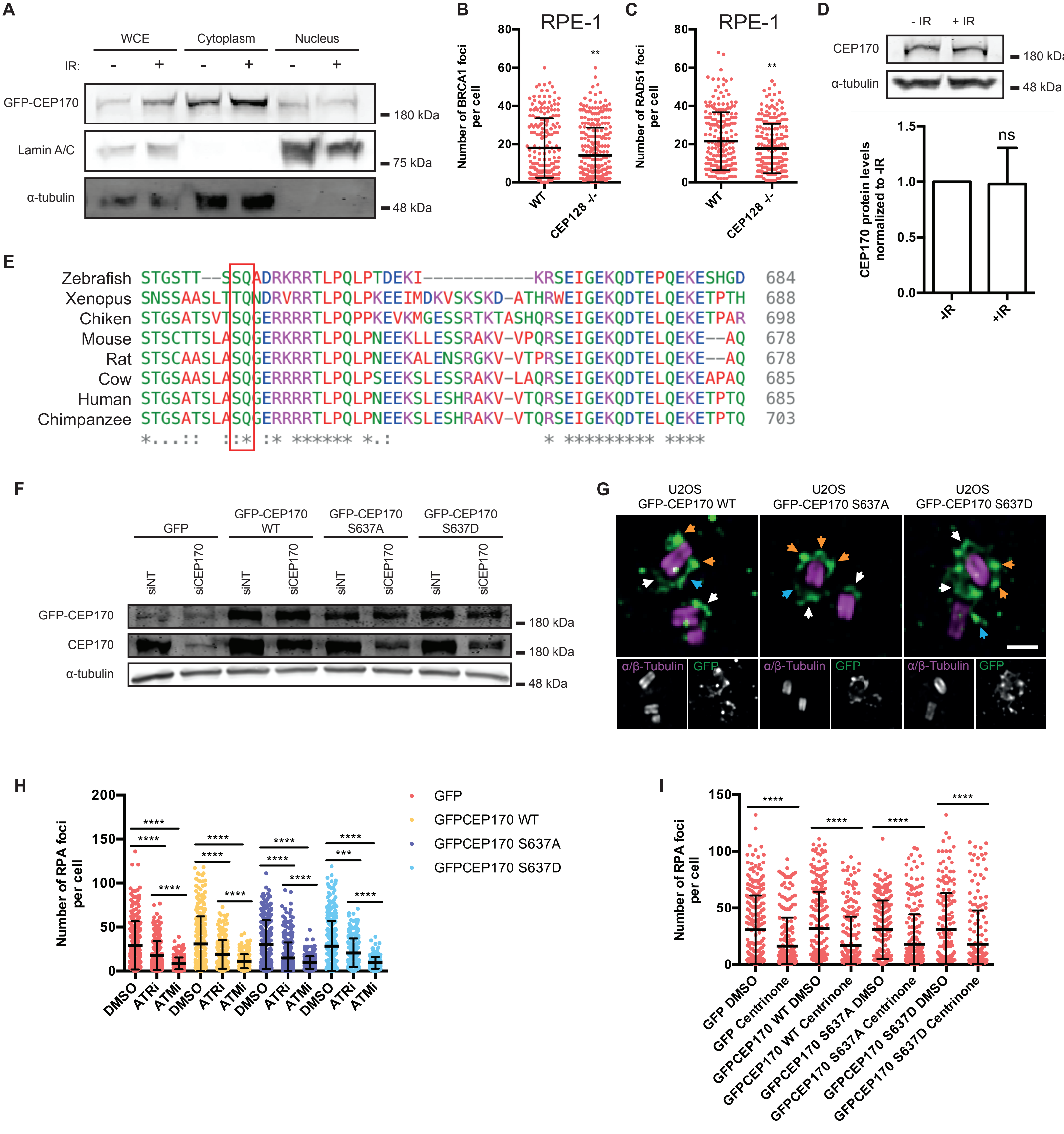


**Rodriguez-Real Figure Expanded View 3**

### Experimental View 4

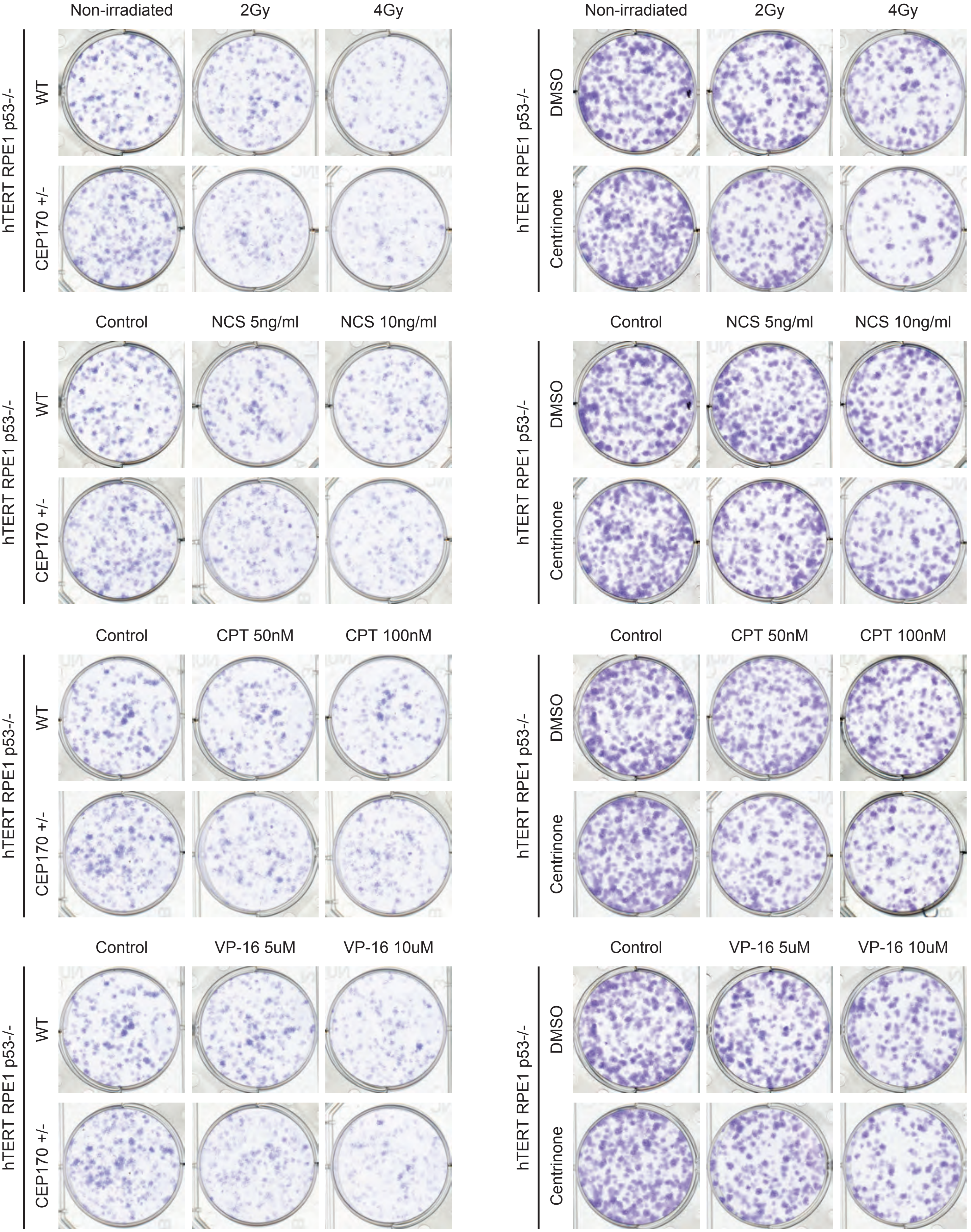


**Rodriguez-Real Figure Expanded View 4**

### Experimental View 5

A

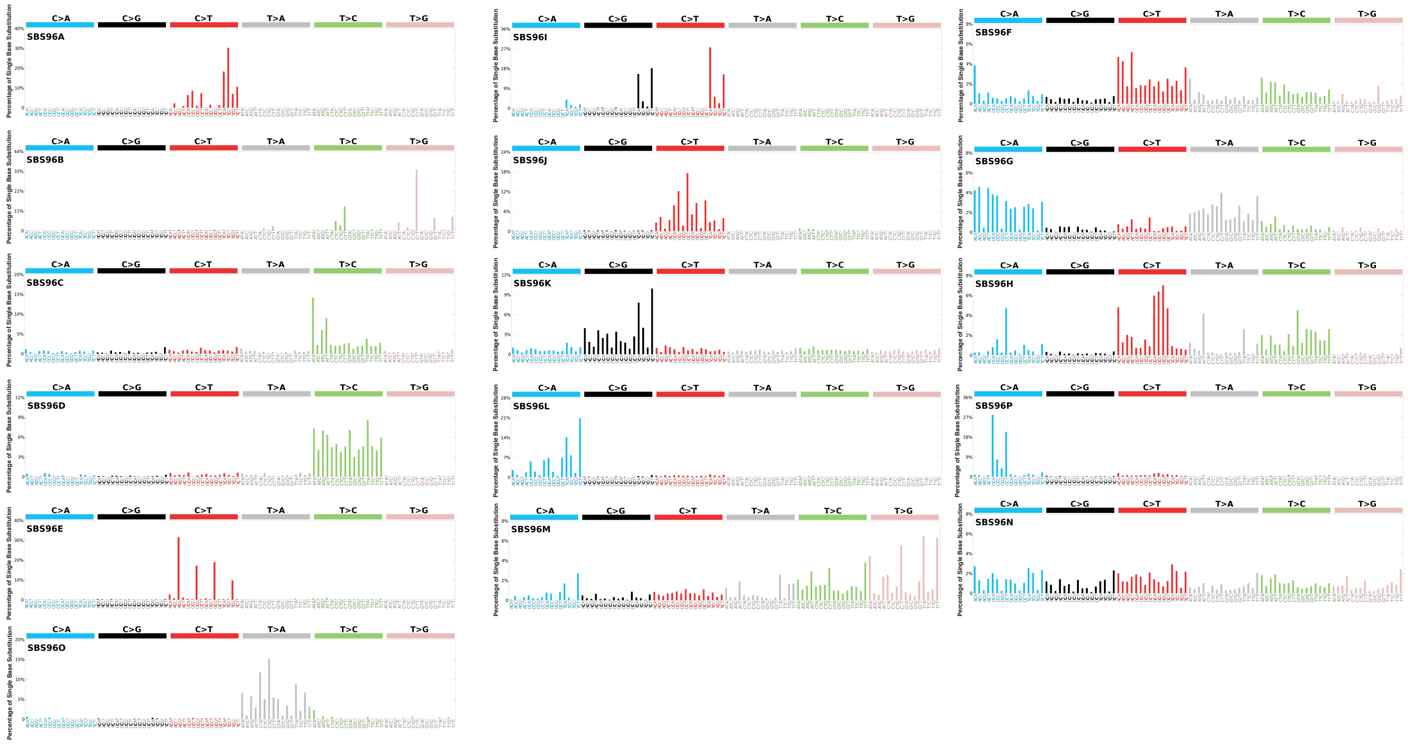


B

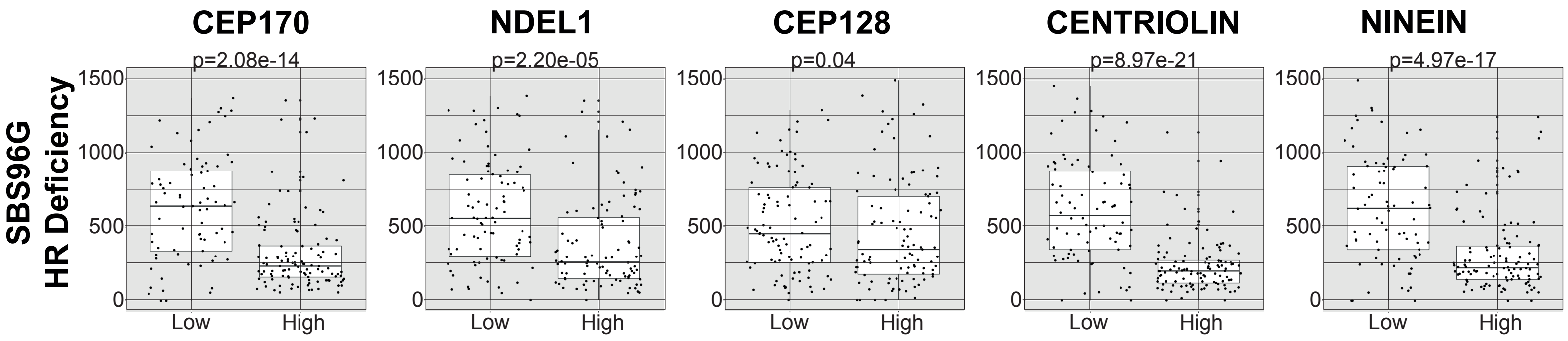


Rodriguez-Real Figure Expanded View 5
