## Appendix for "Centriolar subdistal appendages promote double strand break repair through homologous recombination"

**Rodriguez-Real et al Appendix Table of Contents:**


| | |
|---|---|
| Appendix Figure S1 | page 2-3 |
| Appendix Figure S2 | page 4-6 |

### Appendix Figure S1

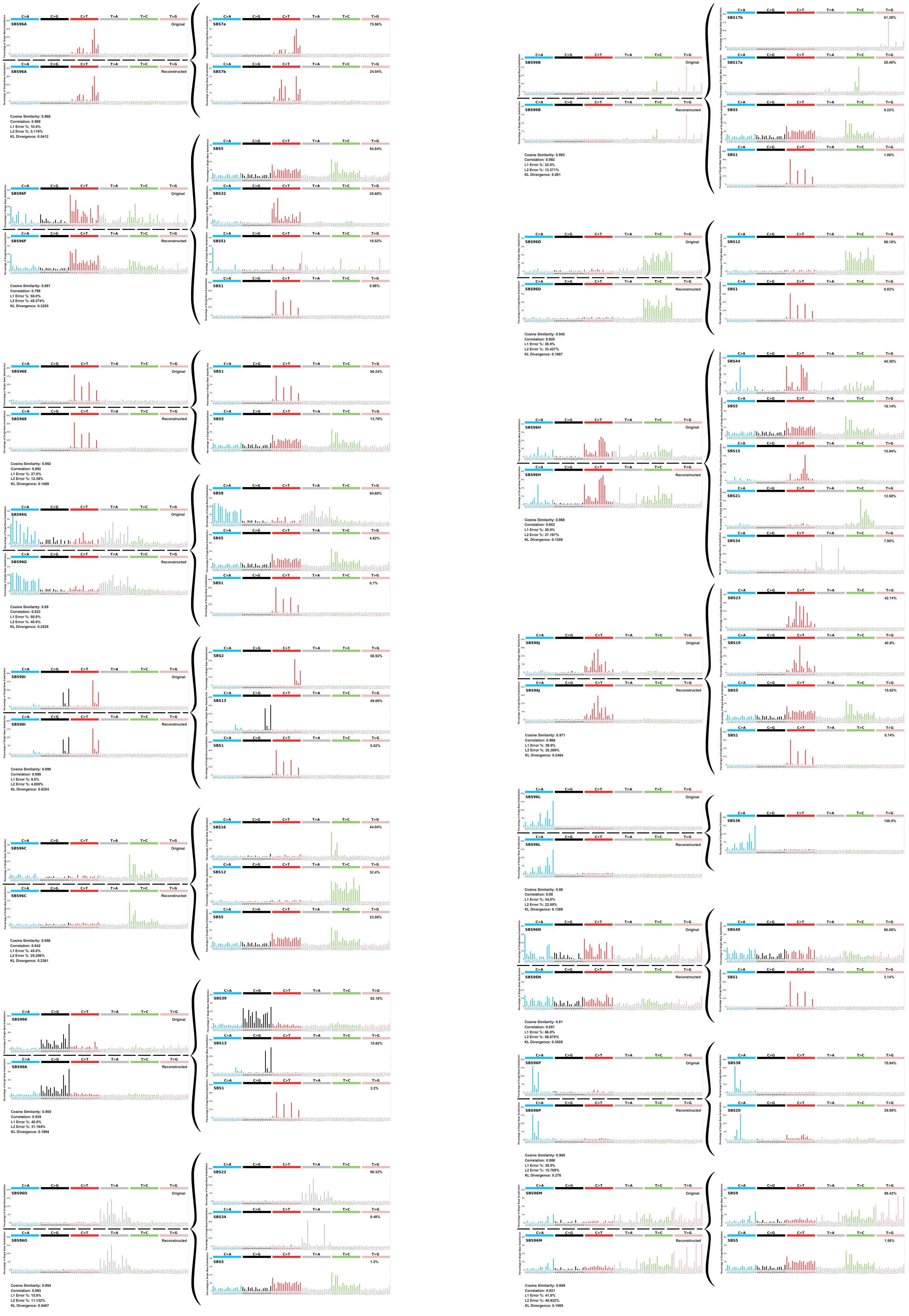

**Figure S1: Comparison of the mutational signatures analyzed with the ones present in COSMIC.** Each signature found in this analysis was compared with the ones described in COSMIC. The combination of specific reference mutational signatures (right side) that when combined (bottom left side "Reconstructed") were similar to the observed signature is shown (top left side "Original"). Statistical parameters of the similarities in each case are shown. The contribution of each reference signature to the reconstructed one is shown.

Appendix Figure S2

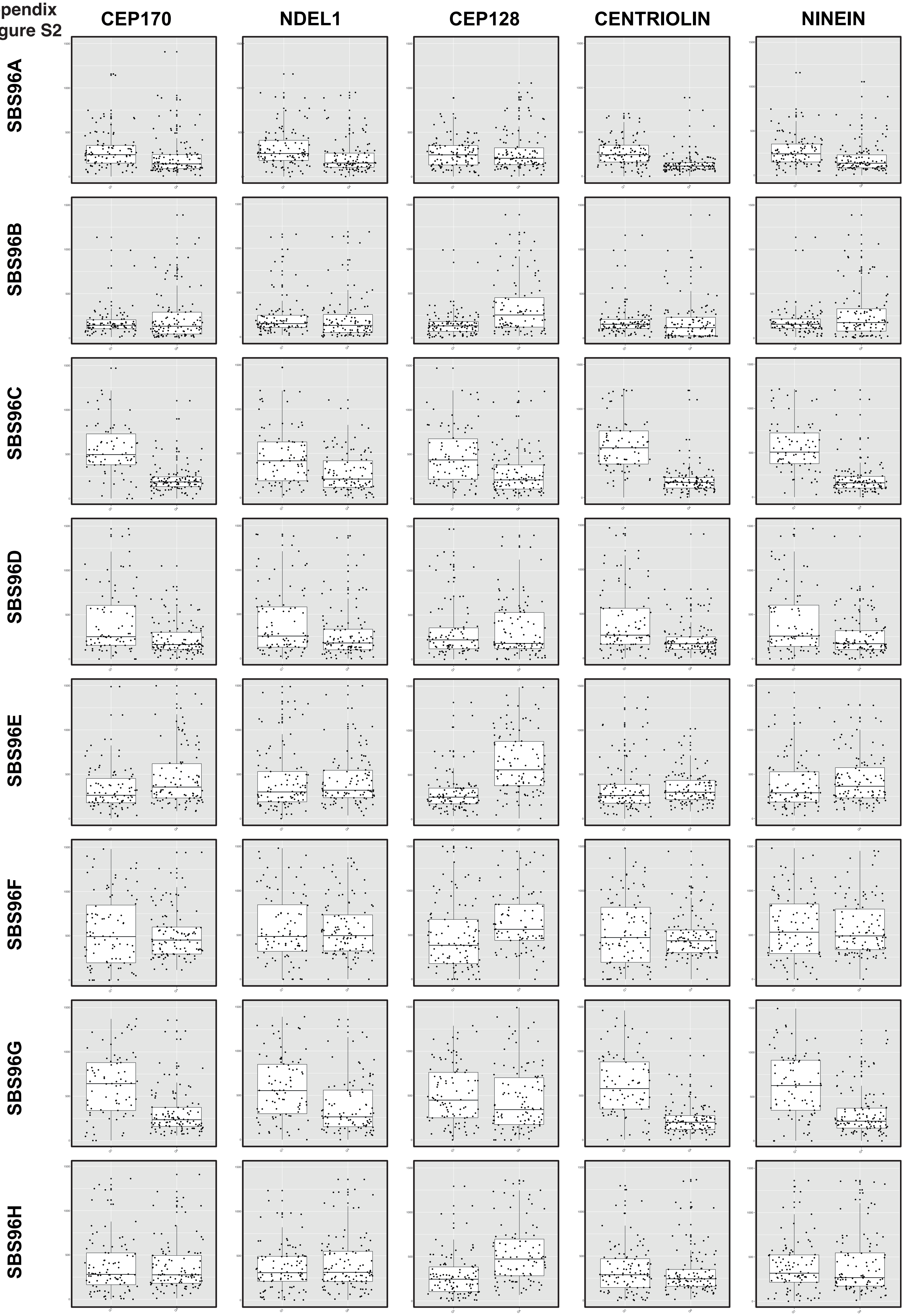

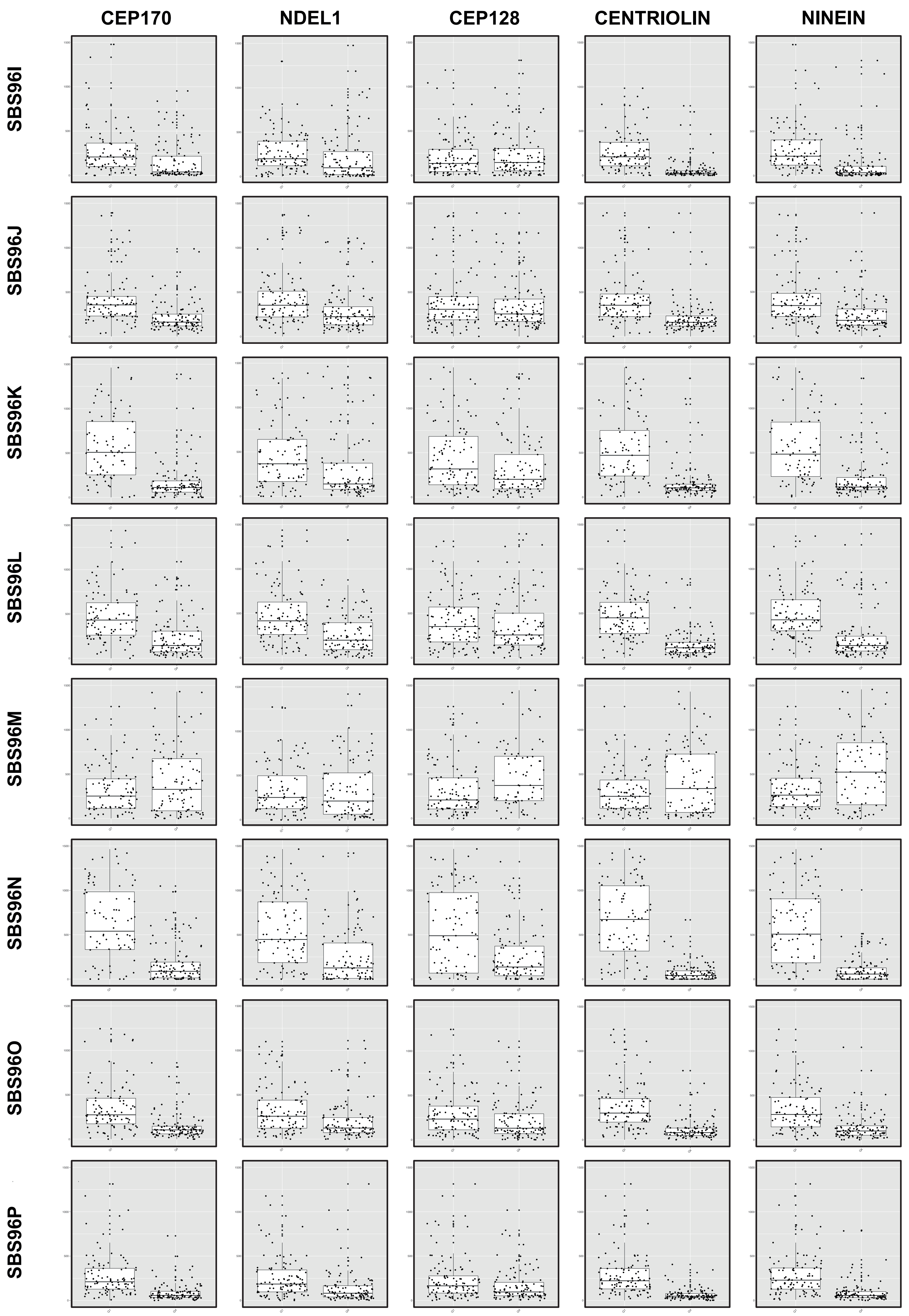
CEP170
NDEL1
CEP128
CENTRIOLIN
NINEIN
SBS96I
SBS96J
SBS96K
SBS96L
SBS96M
SBS96N
SBS96O
SBS96P

**Figure S2. Mutational signatures of subdistal appendage genes.** 16 optimal mutational signatures were analyzed in samples at the lower quartile (low) or at the highest (high) of the indicated genes (on the top). The number of mutations in each sample associated to each signature was plotted. The average is shown. Statistical significance was calculated using a Wilcoxon test, and the p-value is shown on top of the graph. The assigned name of the signature is shown on the side.
